# Respiratory mucosal vaccination of peptide-poloxamine-DNA nanoparticles provides complete protection against lethal SARS-CoV-2 challenge

**DOI:** 10.1101/2022.05.29.493866

**Authors:** Si Sun, Entao Li, Gan Zhao, Jie Tang, Qianfei Zuo, Larry Cai, Chuanfei Xu, Cheng Sui, Yangxue Ou, Chang Liu, Haibo Li, Yuan Ding, Chao Li, Dongshui Lu, Weijun Zhang, Ping Luo, Ping Cheng, Yuwei Gao, Changchun Tu, Bruno Pitard, Joseph Rosenecker, Bin Wang, Yan Liu, Quanming Zou, Shan Guan

## Abstract

The ongoing SARS-CoV-2 pandemic represents a brutal reminder of the continual threat of mucosal infectious diseases. Mucosal immunity may provide robust protection at the predominant sites of SARS-CoV-2 infection. However, it remains unclear whether respiratory mucosal administration of DNA vaccines could confer protective immune responses against SARS-CoV-2 challenge due to the insurmountable barriers posed by the airway. Here, we applied self-assembled peptide-poloxamine nanoparticles with mucus-penetrating properties for pulmonary inoculation of a COVID-19 DNA vaccine (pSpike/PP-sNp). Not only displays the pSpike/PP-sNp superior gene-transfection and favorable biocompatibility in the mouse airway, but pSpike/PP-sNp promotes a tripartite immunity consisting of systemic, cellular and mucosal immune responses that are characterized by mucosal IgA secretion, high levels of neutralizing antibodies, and resident memory phenotype T-cell responses in the lungs of mice. Most importantly, pSpike/PP-sNp completely eliminates SARS-CoV-2 infection in both upper and lower respiratory tracts and enables 100% survival rate of mice following lethal SARS-CoV-2 challenge. Our findings indicate PP-sNp might be a promising platform in mediating DNA vaccines to elicit all-around mucosal immunity against SARS-CoV-2.

## Introduction

The coronavirus disease 2019 (COVID-19) pandemic caused by severe acute respiratory syndrome coronavirus 2 (SARS-CoV-2) has posed a huge and continual threat to the world. Although there are already several authorized COVID-19 vaccines, all these vaccines are inoculated via parenteral route, which predominantly elicit systemic immunity dominated by serum IgG antibodies without conferring mucosal immunity^1^. SARS-CoV-2, which invades the host through the mucosa of the respiratory tract, may seed in the initial reservoir and continue to spread between individuals^2^. Recent studies have demonstrated that respiratory mucosal vaccinated adenoviral vaccines could overcome the drawbacks of parenterally administered counterpart via the stimulation of broad local immune responses in the airways that in turn can block both infection and spread from this reservoir^3–6^. Secretory immunoglobulin A (sIgA) and pulmonary resident memory T cells (TRM) are suggested to be key components in this first-line of defense^7–9^. Compelling evidences have indicated that sIgA in the respiratory mucosa contributed to SARS-CoV-2 neutralization to a greater extent than IgG equivalents^10, 11^, and lung-TRM cells are critical in mediating protection against respiratory pathogens^12^. Since mucosal route of immunization is considered by the research community as the most straightforward approach to induce potent mucosal immunity^13^, safe and efficient delivery of vaccine to the respiratory mucosa would be needed in order to reliably stimulate sIgA secretion and engender protective mucosal immunity against SARS-CoV-2^14^.

Despite the great potential of mucosal immunity, the major advances seen with parenterally administered vaccines (such as adjuvanted subunit antigens, DNA and, more recently, RNA vaccines) have not been translated into licensed mucosal vaccines. There continues to be a dearth of safe and effective respiratory mucosal vaccine platforms despite decades of investigations^15^. Up to date, the only successful approach is virus-based mucosal vaccines. Of the nine mucosal vaccines approved for use in humans (one intranasal and eight oral) all are either live attenuated or whole-cell inactivated vaccines^16^. The dichotomy in parenteral and mucosal approaches is, in part, due to the existence of multiple barriers (e.g., the mucus layer that traps exogenously inhaled substances) and natural defense mechanism (such as mucociliary clearance) keeping the foreign substances out^13^. Viruses have evolved to proficiently overcome these barriers, this explains why the vast majority of investigations achieving protection against SARS-CoV-2 using airway route of immunization exclusively utilized virus-based vaccines, such as adenoviruses^6, 17–19^. Nevertheless, these virus vaccines are often compromised by preexisting immunity and safety issues, some adenovirus-based COVID-19 vaccines may cause cerebral venous sinus thrombosis^20^, rendering the US Food and Drug Administration puts strict limits on this type of COVID-19 vaccines. Therefore, it is difficult to strike a delicate balance between immune efficacy and safety of these virus vaccines. On the other hand, cutting-edge advances and research into the nucleic acid-based vaccine technologies provide alternative options to solve the unmet needs for safe and efficient non-viral based mucosal vaccines.

DNA vaccines, which proved to be helpful in controlling the pandemic^21^, would offer substantial advantages over virus-based vaccines, including safe profiles^22, 23^, ease of manufacture with low costs, and stability at room temperature^21^. However, DNA vaccines generally display limited immunogenicity in the airways because the most commonly used electroporation device is not applicable and consequently require potent delivery systems to overcome the barriers within the respiratory tract^24^. Traditional DNA delivery vectors, such as cationic polymers and even lipid-based formulations that were tested in clinical trials, have turned out to be inefficient owing to the poor airway mucus penetrating properties^25, 26^. Therefore, the pulmonary administration of conventional DNA nanoparticles is unlikely to shuttle DNA cargos efficiently to initiate robust immune responses in the respiratory tract. As revealed by a previous study, the intranasal COVID-19 DNA vaccination in mice only led to a modest induction of local T-cells secreting IFN-γ, without eliciting mucosal sIgA and circulating IgG against SARS-CoV-2 even after prime and two boost doses^27^. To the best of our knowledge, whether a COVID-19 DNA vaccine delivered via the airway could confer protective efficacy against SARS-CoV-2 challenge was still unknown.

In a previous study, we developed a self-assembled peptide-poloxamine nanoparticle (PP-sNp) delivery system that is specifically designed for efficient delivery of plasmid DNA across the mucus layer of the respiratory tract^28^. PP-sNp carrying a *Sleeping Beauty* transposon system successfully enabled the genomic integration of *CFTR* gene in the airway epithelia of cystic fibrosis mice with a safe integration profile^28^. These results suggest that PP-sNp would be promising in mediating DNA vaccines to stimulate potent mucosal immune responses against SARS-CoV-2. To this end, we integrated an intercellular adhesion molecule-1 (ICAM-1/CD54, which is abundantly expressed on the epithelial and immune cells^29^) targeting moiety into the PP-sNp with the purpose of enhancing the specific cellular uptake of pulmonary dendritic cells and airway epithelial cells. These cells have been suggested to play important roles in both initiating potent respiratory immune responses and orchestrating innate immunity to maintain normal airway architecture^30, 31^. A DNA vaccine encoding the wild-type spike protein of SARS-CoV-2 (pSpike, which has been tested in clinical trials^22^) was formulated in PP-sNp (pSpike/PP-sNp) for mucosal vaccine applications. Three vaccine doses of pSpike/PP-sNp via respiratory tract induced comprehensive and broad protective immune responses, including mucosal immunity (SARS-CoV-2 specific sIgA in the airways as well as lung-TRM cells) and systemic immunity (neutralizing serum IgG). All these merits conferred virtually full protection of vaccinated mice against lethal SARS-CoV-2 challenge and completely removed the virus in both the turbinates and lungs of mice. Thus, our study may serve as the first proof-of-concept revealing that respiratory mucosal immunization of DNA vaccines holds the potential to elicit robust protective immunity against SARS-CoV-2 infection in the upper and lower respiratory tracts.

## Results

### The characterization and in vitro investigation of peptide-poloxamine nanoparticle (PP-sNp) containing plasmid DNA

The PP-sNp formulation is formed by a simple self-assembly of multi-modular peptide (Supplementary Table 1), poloxamine 704 (a block copolymer proven to be capable of mediating efficient DNA transfection in the airways to a level that is significantly better than “gold-standard” lipid-based GL67A formulation that utilized in clinical trials^26, 32, 33^) and pDNA components (Fig. 1a). We engineered small and monodisperse PP-sNp nanoparticles with negative charge (ζ potential: −34.7±0.5 mV) via an optimized procedure in order to deal with the small volumes and high concentrations required in vivo (Fig. 1b, Supplementary Table 2). The electron microscopy showed that PP-sNp had a spherical morphology with uniform size distribution (Fig. 1c and 1d). PP-sNp can be stored at room or high temperatures and remains stable (Supplementary Fig. 1). We then evaluated the uptake efficiency of PP-sNp in DC2.4 and MH-S cell lines, the results indicate an enhanced cellular uptake of PP-sNp containing Cy5 labelled pDNA (Cy5-pDNA) by both cell lines compared to the 25 kDa branched polyethyleneimine (PEI, a representative polymer based delivery vehicle in DNA mucosal vaccines^34–36^)-based and naked Cy5-pDNA-based controls (Fig. 1e, Supplementary Fig. 2). Confocal microscopy revealed that the majority of Cy5-fluorescence was observed in the nucleus of Cy5-pDNA/PP-sNp transfected DC2.4 cells with dispersed patterns, while the Cy5-fluorescence signal within the nucleus of cells transfected by Cy5-pDNA/PEI and naked Cy5-pDNA was very limited (Fig. 1f). Afterwards, pDNA encoding firefly luciferase (pFLuc) was applied as a reporter to evaluate their transfection efficiency. As shown in Fig. 1g and Supplementary Fig. 3, PP-sNp displayed the most efficient transfection in several cell lines (DC2.4, Calu-3, 16HBE and BEAS-2B) compared to control formulations, such as lipofectamine2000 (lipo2000)-based lipoplex, PEI-based polyplex and naked-pFLuc. Meanwhile, we evaluated the cytotoxicity of these formulations and found PP-sNp based formulation did not provoke significant cytotoxicity on the viability of different cell lines, whereas Lipo2000-based and PEI-based counterparts did (Fig. 1h). Additionally, multiple particle tracking (MPT) assays were performed to evaluate the mucus penetrating ability of PP-sNp. The trajectories of the PP-sNp-based and PEI-based particle motions were captured (Supplementary Video 1 and 2), and representative trajectories were mapped in Fig. 1i. PEI nanoparticles were almost trapped by the mucus network. In contrast, PP-sNp was able to move freely in a large area, displaying a better diffusion pattern. The mean square displacement (〈MSD〉) of PP-sNp was about 1000-fold higher than PEI counterpart at 10 s (Fig. 1j). Consistently, the effective diffusivities (Deff) of PP-sNp were significantly higher than that of PEI (Fig. 1k). The distinct movement patterns suggest that PP-sNp hold the capability of efficient mucus penetrating.

**Fig. 1.**
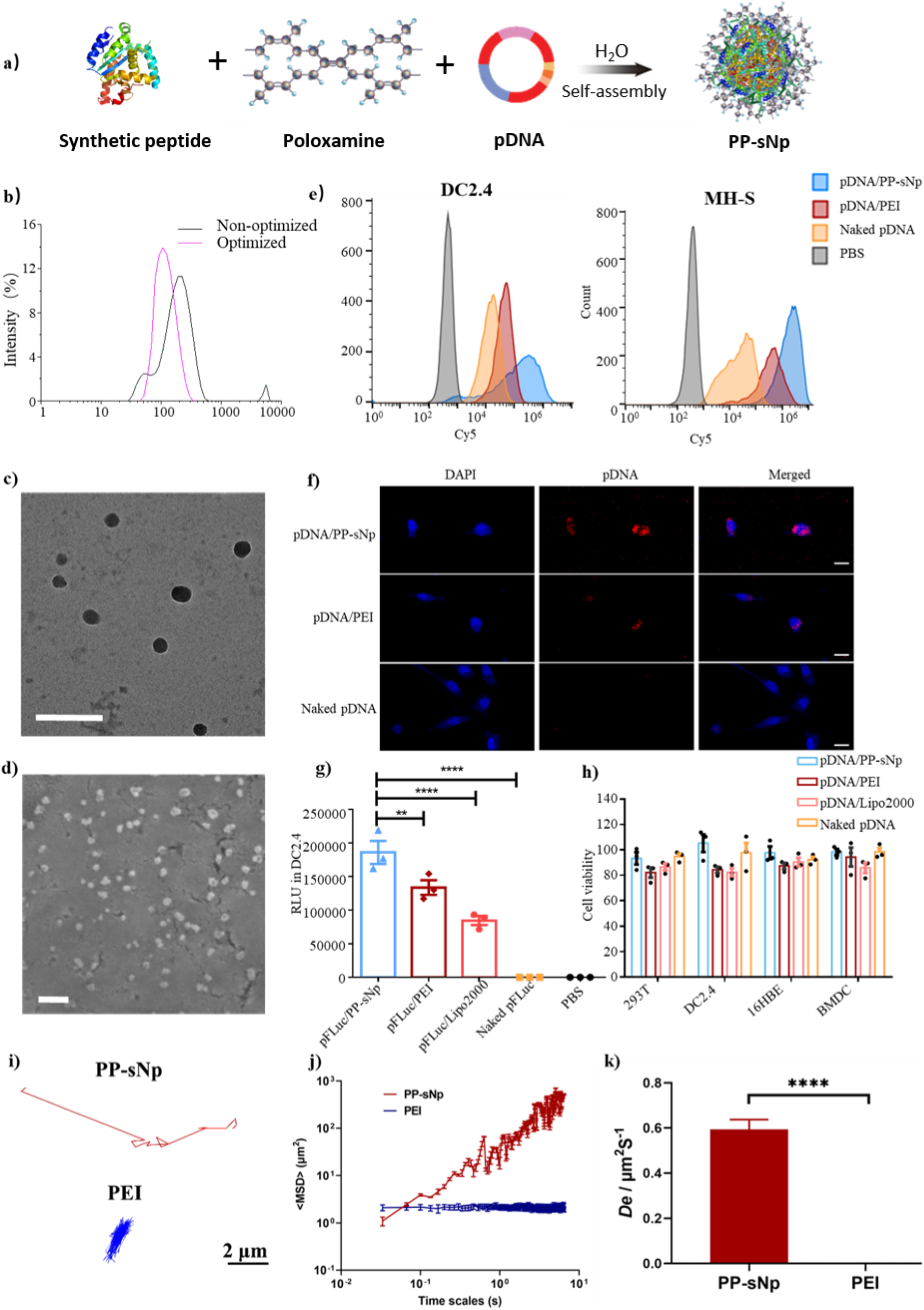
Characterization and in vitro evaluation of pDNA/PP-sNp. **a**) Schematic showing the formation of pDNA/PP-sNp through a self-assembly process in aqueous solution. **b**) The dynamic laser scattering results measured for particle sizes of pDNA/PP-sNp prepared by different methods. **c**) The transmission electron microscopy (TEM) micrograph and **d**) the scanning electron microscope (SEM) micrograph of pDNA/PP-sNp. Scale bar: 200 nm. **e**) Cellular uptake of Cy5-pDNA/PP-sNp, Cy5-pDNA/PEI and naked Cy5-pDNA in DC2.4 and MH-S cells. Cy5-fluorescence intensity within the cells was measured by flow cytometry. **f**) Subcellular fate of the Cy5-pDNA/PP-sNp, Cy5-pDNA/PEI and naked Cy5-pDNA after 4 h incubation with DC2.4 cells detected by confocal laser scanning microscopy. Blue channel, nuclei stained by DAPI; Red channel, Cy5-pDNA; Merged, combination of the aforementioned channels. Scale bars: 20 μm. **g**) Transfection efficiency of pFLuc/PP-sNp in DC2.4 cells. DC2.4 cells were incubated with pFLuc/PP-sNp, pFLuc/PEI, pFLuc/lipo2000 or naked-pFLuc for 4 h, then subjected to detection of bioluminescence in cell lysate 48 h post transfection. Statistical significance was calculated by one-way ANOVA with Dunnett’s multiple comparisons tests (\*\**P*<0.01, \*\*\*\**P*<0.0001). **h**) CCK-8 cell viability assay of pDNA/PP-sNp, pDNA/PEI, pDNA/lipo2000 or naked-pDNA after 4 h incubation with different cell lines (all formulations were prepared under conditions showing the highest transgene expression). Data in **g** and **h** represent mean ± SEM (n = 3 independent experiments). Multiple particle tracking (MPT) studies were carried out to investigate the movement of PP-sNp in mucus mimicking gel, the motion of PEI nanoparticles was quantified for comparison as a control. **i**) Representative trajectories for particles in mucin solution (3%, w/v) during the 15-s movies. **j**) The mean square displacements (<MSD>) of individual nanoparticle as a function of the time scale. **k**) The effective diffusivities (Deff) of individual nanoparticle (n > 100 nanoparticles per experiment). Data in **j** and **k** represent mean ± SEM of three independent experiments. A two-tailed unpaired *t*-test was used to determine the significance of the indicated comparisons (\*\*\*\**P*< 0.0001).

### In vivo evaluation of the transfection properties and biocompatibility of PP-sNp

Intratracheal application of PP-sNp led to strong bioluminescence signals in the lungs of mice 24 h later, the signal reached peak 48 h post-dosing and declined to the background level 7 days afterwards (Fig. 2a and Supplementary Fig. 4). Bioluminescence signals from live animals and excised organs reveal intratracheally administered PP-sNp exhibited superior efficacy compared to pFLuc/PEI or naked-pFLuc counterparts, and the lung was the most abundant fLuc-expressing site and no signals were detected in other organs (e.g., heart, live, spleen, kidneys and small intestinal) after the PP-sNp transfection (Fig. 2b). Based on these results, we further evaluated the capability of PP-sNp in intratracheal delivery of a plasmid encoding the full-length spike protein of SARS-CoV-2 (pSpike)^22^. Considerable levels of spike gene-specific mRNA were detected in the lungs of mice transfected by pSpike/PP-sNp compared to pSpike/PEI or naked-pSpike counterparts (Fig. 2c). Histopathological analysis demonstrated that the lungs and other major organs (including heart, liver, spleen and kidney) from pSpike/PP-sNp treated mice were indistinguishable from naked-pSpike and phosphate-buffered saline (PBS) controls, without significant histopathological signs of inflammation (Supplementary Fig. 5). In contrast, administration of pSpike/PEI counterpart resulted in signs of interstitial oedema, damage to the epithelial barrier, and overt infiltration of inflammatory cells in the lung (Supplementary Fig. 5), which is in consistent with previous findings indicating that the highly positively charged and non-degradable nature of PEI based formulation tends to provoke the activation of genes with apoptosis, stress responses, and oncogenesis^37^. PEI-based formulation was excluded in the following studies due to its poor transfection efficiency and toxic profiles. Although there was a lack of overt lung inflammation observed with histology, pSpike/PP-sNp may rapidly and robustly, but only transiently, activate innate immunity. As shown in Fig. 2d, tumor necrosis factor-α (TNF-α) and interleukin-6 (IL-6) expression within bronchoalveolar lavage fluid (BALF) and supernatants of lung homogenates peaked during 1-2 days after administration then immediately resolved to background levels. However, no significant increase of cytokines was found in the spleen-based and serum-based counterparts (Supplementary Fig. 6).

**Fig. 2.**
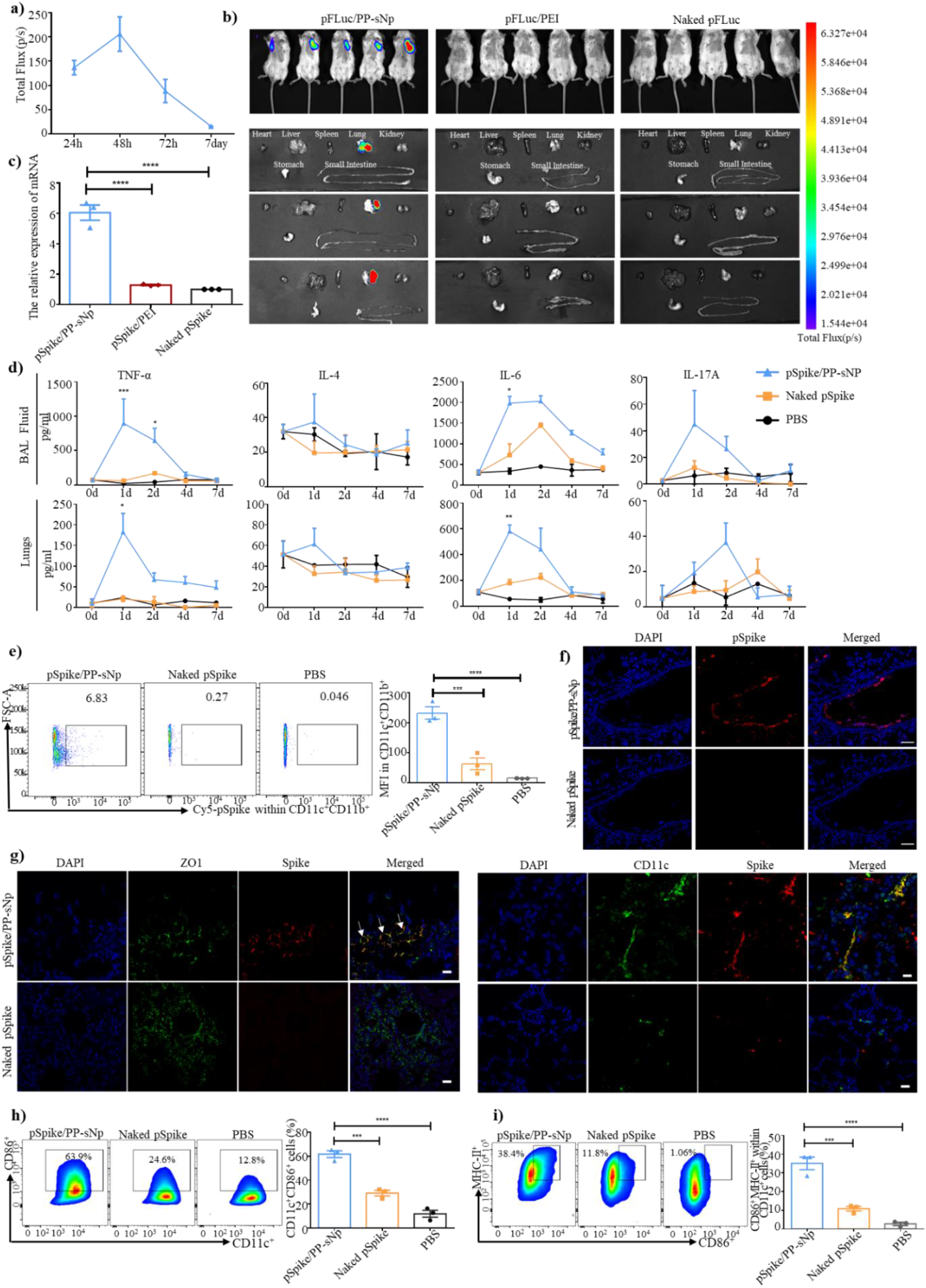
In vivo transfection profiles and biocompatibility of pDNA/PP-sNp. **a**) Kinetics of transgene expression mediated by intratracheally administered pFLuc/PP-sNp over time in mice. Data represent mean ± SEM (n = 5 biologically independent samples). **b**) Images of in vivo bioluminescence induced by pFLuc/PP-sNP, pFLuc/PEI and naked-pFLuc 48 h after intratracheal dosing in mice. **c**) The expression of the SARS-CoV-2 spike gene specific-mRNA in lung of mice intratracheally administered pSpike/PP-sNp, pSpike/PEI and naked-pSpike, measured by RT-qPCR 48 h after transfection. **d**) Supernatants of lung homogenates and BALF samples were collected at indicated time points and were analyzed for cytokine levels via enzyme-linked immunosorbent assay (ELISA). Data represent mean ± SEM (n = 3 biologically independent samples). **e**) In vivo DC uptake of pSpike/PP-sNp and naked-pSpike 15 h following intratracheal administration. Mice were sacrificed to detect the fluorescence intensity of Cy5 in pulmonary DC cells via flow cytometry. **f**) Representative confocal images demonstrating the mucus-penetration properties of Cy5-pSpike/PP-sNp and naked Cy5-pSpike through airway mucus of mice 15 h after the administration. Blue channel, nuclei stained by DAPI; Red channel, Cy5-pSpike Scale bars: 50 μm. **g**) Immunohistochemical staining of the spike protein (red channel) and CD11c (green channel) or ZOl (zonula occludes protein 1, which indicates the epithelium, green channel) in lung of mice treated by pSpike/PP-sNp and naked-pSpike via pulmonary route. Scale bars: 25 μm. **h**) Expression of CD11c^+^CD86^+^ or **i**) CD86^+^MHC-II^+^ (gated on CD11c^+^) on BMDCs after 24 h incubation with pSpike/PP-sNp, naked-pSpike and phosphate buffered saline (PBS) (n = 3 biologically independent samples). Each symbol in the bar chart of **c**, **e**, **h**, and **i** represents one sample from a biologically independent mouse. Data in **c**, **e**, **h** and **i** are shown as mean ± SEM. One-way ANOVA with Dunnett’s multiple comparisons tests was used to determine significance (\**P*<0.05, \*\**P*<0.01, \*\*\**P*<0.001, \*\*\*\**P*<0.0001).

Mature DCs are powerful antigen-presenting cells (APCs) and play key roles in initiating antigen-specific immune responses. We therefore evaluated whether the pulmonary DCs (CD11c^+^CD11b^+^) can efficiently uptake the pSpike/PP-sNp. Comparative analysis showed that 6.15±0.65% of pulmonary DCs internalized Cy5-pSpike/PP-sNp with a mean fluorescence intensity (MFI) of 232.0±20.60, which was significantly higher than naked Cy5-pSpike counterpart with a 1.23±0.64% uptake rate (MFI: 62.5±19.90) (Fig. 2e, Supplementary Fig. 7). Meanwhile, we confirmed a deep penetration and widespread distribution of the intratracheally administered Cy5-pSpike/PP-sNp in the mucus-covered respiratory tract in vivo (Fig. 2f). On the contrary, identically administered naked-pSpike was almost undetectable (Fig. 2f), demonstrating the superior in vivo mucus penetrating ability of PP-sNp. Immunofluorescence sections of airway region suggest that considerable pSpike expression mediated by PP-sNp could be clearly observed in pulmonary DCs and airway epithelial cells (AECs) with a widespread distribution pattern, while the spike protein signal within naked-pSpike treated group was very limited and neither co-localize with pulmonary DCs nor AECs (Fig. 2g). To our surprise, we could not detect transgene expression mediated by pSpike/PP-sNp in alveolar macrophages (Supplementary Fig. 8), which are representative airway APCs being supposed to engulf most of the exogenous antigens^38^. We further examined the ability of pSpike/PP-sNp in promoting DC maturation. Compared to naked-pSpike, pSpike/PP-sNp could induce two-fold expressions of CD11c^+^CD86^+^ on bone marrow derived cells (BMDCs) (Fig. 2h and Supplementary Fig. 9). The expressions of CD86^+^MHC-II^+^ on CD11c^+^ cells were significantly elevated by more than 300% in pSpike/PP-sNp treated group compared to naked-pSpike control (Fig. 2i and Supplementary Fig. 9).

### pSpike/PP-sNp induces robust mucosal and humoral immunity

To determine the immunogenicity of mucosal vaccination, mice were immunized via intratracheal route with pSpike/PP-sNp, naked-pSpike, an empty vector pVax loaded PP-sNp (pVax/PP-sNp, served as Mock control) or PBS (Fig. 3a). We observed the body weight of mice treated by pSpike/PP-sNp and naked-pSpike transiently decreased during 2-3 days post the prime and 1^st^ boost dosing, which progressively recovered to the levels displayed by other control groups (Fig. 3b). We also confirmed the necessity of the 2^nd^ boost dose by detection of IgG in serum and sIgA in BALF at pre-determined time points (Supplementary Fig. 10). Intratracheal immunization of pSpike/PP-sNp, but neither pVax/PP-sNp, naked-pSpike nor PBS controls, induced high levels of spike-specific IgG antibodies in serum samples (Fig. 3c). The highest IgG titer within serum from pSpike/PP-sNp group reached 1/51200 on day 35 (Fig. 3d). The titer ratio of IgG2a/IgG1 implies that pSpike/PP-sNp tends to induce a Type 1 T helper cell (Th1)-biased immune response (Supplementary Fig. 11). Most importantly, only pSpike/PP-sNp induced high levels of spike-specific sIgA antibody in BALF with most samples could reach an endpoint titer of 1/128 (Fig. 3e and Fig. 3f). Furthermore, distal mucosal immune responses were also induced by pSpike/PP-sNp as it proved by the presence of SARS-CoV-2-specific sIgA antibody in vaginal lavage fluid (Supplementary Fig. 12). The neutralization titer (ND_50_) of serum and BALF samples from pSpike/PP-sNp group approached ~1/1875 (Fig. 3g) and ~1/73 (Fig. 3h), respectively. Whereas no neutralizing antibodies (NAb) could be detected in samples from other groups (naked-pSpike, pVax/PP-sNp and PBS). Besides, we successfully detected antibody-secreting plasma cells producing IgA or IgG against SARS-CoV-2 in the spleen after intratracheal immunization with pSpike/PP-sNp (Supplementary Fig. 13).

**Fig. 3.**
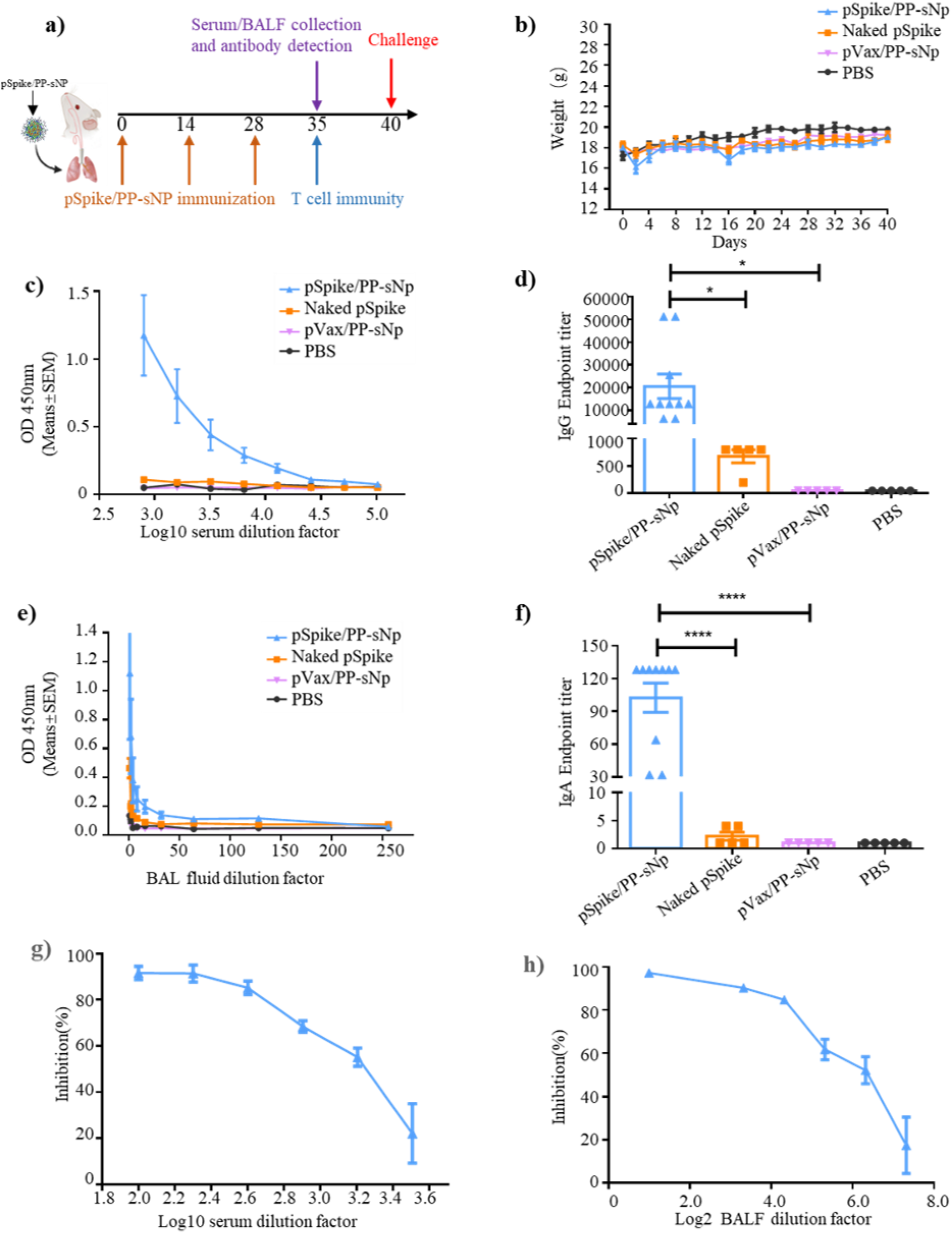
Humoral immune responses after intratracheal immunization of pSpike/PP-sNp in mice. **a**) Schematic diagram of immunization, sample collection and challenge schedule. Mice were immunized on day 0 and boosted with the same dose on day 14 and day 28, respectively. **b**) Animal weights were recorded during the whole period of immunization. Data represent mean ± SEM (n= 10 mice/group). **c**) OD450 nm values of SARS-CoV-2 spike (S1 + S2) protein-specific IgG in serial diluted serum samples collected 35 days after initial vaccination. Data represent mean ± SEM (n= 10 biologically independent samples). **d**) The SARS-CoV-2 spike (S1 + S2) protein-specific IgG antibody titer in serum samples collected 35 days after initial vaccination. Each symbol represents one sample from a biologically independent mouse. **e**) OD450 nm values of SARS-CoV-2 spike (S1 + S2) protein-specific sIgA in serial diluted BALF samples collected 35 days after priming. Data represent mean ± SEM (n= 10 biologically independent samples). **f**) The SARS-CoV-2 spike (S1 + S2) protein-specific sIgA antibody titer in BALF samples collected 35 days after initial vaccination. Each symbol represents one sample from a biologically independent mouse. Pseudovirus neutralizing antibody titer in **g**) serum samples and **h**) BALF samples collected 35 days after priming. Data in **g** and **h** are shown as mean ± SEM of samples collected from three mice. One-way ANOVA with Dunnett’s post-hoc test was used to determine significance within d and f (\**P*< 0.05, \*\**P*<0.01, \*\*\**P*<0.001, \*\*\*\**P*< 0.0001).

### pSpike/PP-sNp elicits potent T cell responses and memory-biased immunity

In order to investigate cellular immune responses activated via mucosal immunization, we assessed the production of interferon-γ (IFN-γ) and interleukin-4 (IL-4) in pulmonary lymphocytes or splenocytes using ELISpot assay. A significantly higher level of IFN-γ and IL-4 secretion was detected in pulmonary lymphocytes from pSpike/PP-sNp group, while negligible levels of both cytokines were detected in samples from other groups (Fig. 4a and 4b). A similar trend was also observed in splenocyte samples (Supplementary Fig. 14-15). We next evaluated the intracellular IFN-γ and IL-4 production within CD4^+^ and CD8^+^ T cells after *ex vivo* re-stimulation. Flow cytometry analysis showed that pSpike/PP-sNp led to a significant secretion of IFN-γ^+^ by CD4^+^ (1.412±0.077)% (Fig. 4c) and CD8^+^ T cells (1.212±0.102)% (Fig. 4d) within pulmonary lymphocytes. However, there was no significant difference in IL-4 secretion by CD4^+^ T cells between pSpike/PP-sNp immunized mice and naked-pSpike treated ones (Fig. 4e). We also failed to detect significant secretion of IFN-γ or IL-4 in splenic CD8^+^ T and CD4^+^ T cells (Supplementary Fig. 16-17). These data reveal a comprehensive SARS-CoV-2-specific Th1 and cytotoxic T cells activation in the lung of pSpike/PP-sNp-treated mice. To determine whether memory T cell responses within pulmonary mucosal sites were elicited by pSpike/PP-sNp, the effector memory T (T_EM_) cells or central memory T (T_CM_) cells, identified as CD44^hi^CD62L^lo^ and CD44^hi^CD62L^hi^, were detected by flow cytometry. We found both CD4^+^ T_EM_ and CD8^+^ T_EM_ cells in lung, but not splenic T_EM_ cells or T_CM_ cells, were significantly activated via the immunization of pSpike/PP-sNp (Fig. 4f and 4g, Supplementary Fig. 18-19). Recent studies have suggested that tissue-resident memory T (T_RM_) cells play crucial roles in maintaining long-term protective immunity against mucosal pathogens^39, 40^. We investigated the expression of the tissue-retention markers CD69 and CD103 on total CD4^+^ or CD8^+^ T cells isolated from the lung. As shown in Fig. 4h, CD4^+^CD69^+^ cells and CD8^+^CD69^+^ cells in pSpike/PP-sNp immunized group displayed 5.6- and 3.3-fold increase than naked-pSpike counterpart, respectively. Although the activation of CD103 maker was relatively limited, the presence of CD4^+^CD69^+^CD103^+^ cells and CD8^+^CD69^+^CD103^+^ cells within pSpike/PP-sNp immunized group could be identified and was significantly higher than other controls (Fig. 4i). These results denote the tremendous potential of pSpike/PP-sNp in inducing comprehensive mucosal immune responses and immune memory to impart a powerful anti-SARS-CoV-2 protection.

**Fig. 4.**
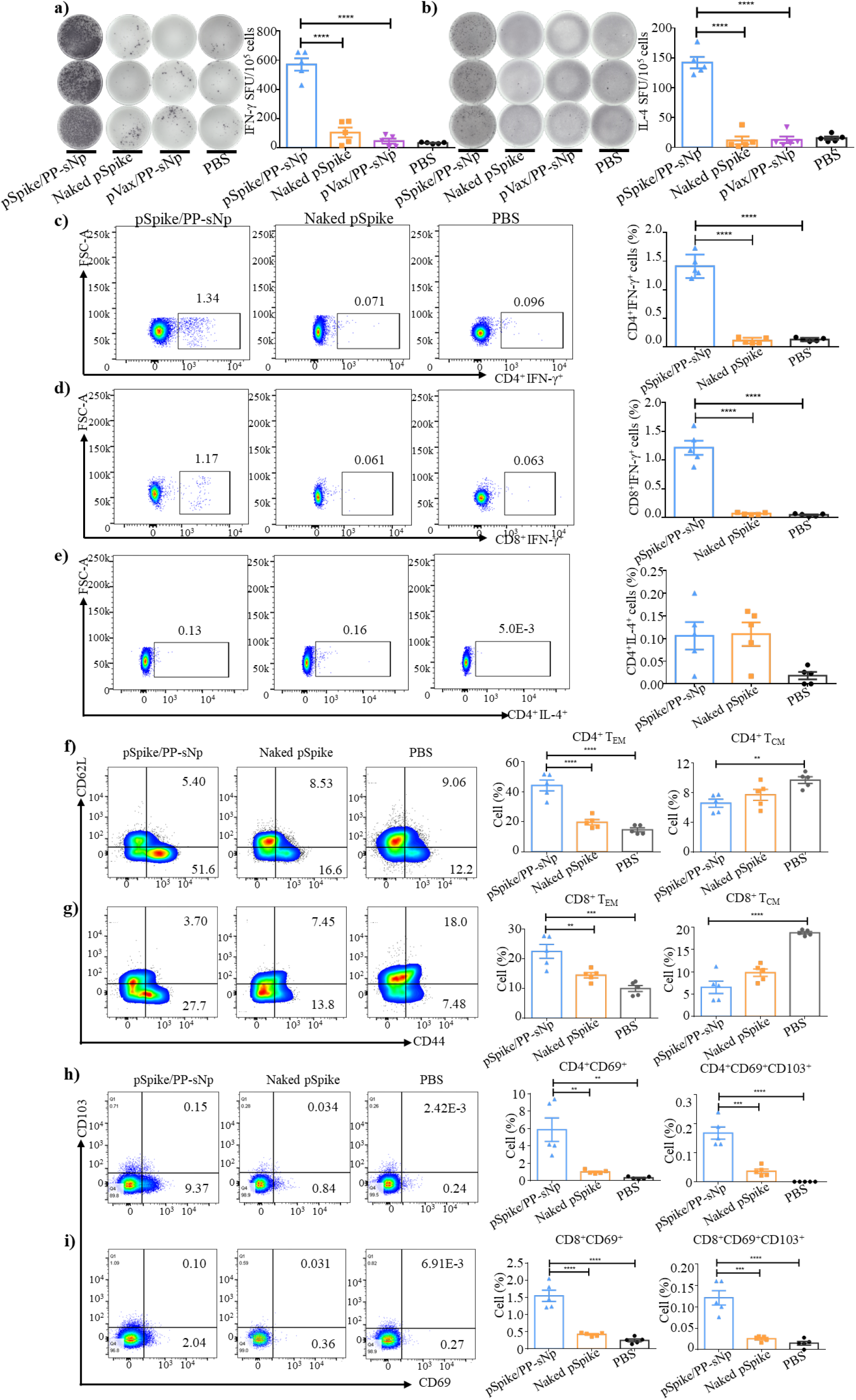
SARS-CoV-2-specific T cell immune responses and memory-biased immunity in pSpike/PP-sNP vaccinated mice. Enzyme-linked immunospot (ELISpot) analyses of **a**) IFN-γ and **b**) IL-4 spot-forming cells in pulmonary lymphocyte after re-stimulation with peptide pools of 14-mer overlapping peptides spanning the SARS-CoV-2 receptor binding domain (RBD) region. **c**) CD4^+^ T cells and **d**) CD8^+^ T cells in the lung were assayed for IFN-γ^+^ expression by flow cytometry after re-stimulation with the SARS-CoV-2 RBD peptide pool. **e**) CD4^+^ T cells in the lung were analyzed for IL-4^+^ expression via flow cytometry in the same way as describe above. **f**) Percentage of CD4^+^ effector memory T cells (TEM) co-expressing CD44^hi^ and CD62L^lo^ and central memory T cells (TCM) co-expressing CD44^hi^ and CD62L^hi^ in the lung of mice 35 days after priming. **g**) Percentage of CD8^+^ T_EM_ and T_CM_ in lung of mice 35 days after priming. **h**) Percentage of CD4^+^ T_RM_ and **i**) CD8^+^ T_RM_ in lung of mice 35 days after initial vaccination. Data in **a-i** represent mean ± SEM (n=5 biologically independent samples). One-way ANOVA with Dunnett’s post-hoc test was used to determine significance (\*\**P*< 0.01, \*\*\**P*< 0.001, \*\*\*\**P*< 0.0001).

### Immunization with pSpike/PP-sNp completely prevents lethal SARS-CoV-2 infection in the upper and lower respiratory tracts

The data described above prompted us to confirm if sufficient protection against SARS-CoV-2 could be achieved via mucosal immunization of pSpike/PP-sNp. To this end, a SARS-CoV-2 C57MA14 strain (which causes severe respiratory symptoms and mortality in mice) was applied in the challenge study. The pSpike/PP-sNp vaccine demonstrated remarkable protective efficacy as evidenced by an 100% survival rate at the predetermined end (14 days post-challenge), while all mice from control groups were deceased during 4-7 days after challenge (Fig. 5a). The body weights of pSpike/PP-sNp vaccinated mice decreased slightly (<10%) in the first 3 days post-challenge but gradually recovered to a level displayed by the control group (without challenge). In contrast, pVax/PP-sNp and PBS groups led to a dramatic decrease in body weight until 6 days after challenge when all animals died (Fig. 5b). Some mice were randomly euthanized to analysis the viral RNA loads in the airways 3 days post-challenge. Significant lower levels of viral RNA were detected in the turbinates (Fig. 5c and 5e) and the lungs (Fig. 5d and 5f) of pSpike/PP-sNp vaccinated mice compared to control counterparts. Moreover, the viral RNA loads in turbinates and lungs of pSpike/PP-sNp treated mice decreased with time and were almost undetectable 14 days post-challenge (Fig. 5g and 5h), indicating pSpike/PP-sNp not only prevents virus infection but eventually eliminates the virus in respiratory tract completely. Additionally, lung sections were subjected to an immunohistochemistry assay aiming to explore the replication of SARS-CoV-2 by detecting its SARS-CoV-2 N protein expression in the lung 3 days post-challenge. Less SARS-CoV-2 N protein in lung sections from pSpike/PP-sNp immunized mice was detected (Fig. 5i). On the other hand, mice treated by pVax/PP-sNp or PBS not only been detected with obvious SARS-CoV-2 N protein (Fig. 5i), but also developed typical lung lesions characterized by denatured epithelial tissues, thickened alveolar septa, and activated inflammatory cell infiltration according to the histopathological assays (Fig. 5j). Whereas the pathological changes significantly alleviated in lung sections from pSpike/PP-sNp vaccinated mice (Fig. 5j). We further evaluated the potential roles of NAb and T_RM_ cells in the process of eliminating SARS-CoV-2. As depicted in Supplementary Fig. 20, significant numbers of CD4^+^T_RM_ cells were identified exclusively in pulmonary sections of pSpike/PP-sNp treated mice after challenge, and the NAb titers against SARS-CoV-2 C57MA14 strain in serum of pSpike/PP-sNp immunized mice were elevated more than 5 times 14 days after challenge (Supplementary Fig. 21), implying NAb and T_RM_ mediated protective effects were probably involved in the protection of mice from lung lesions and the complete elimination of infected SARS-CoV-2 in the whole respiratory tract.

**Fig. 5.**
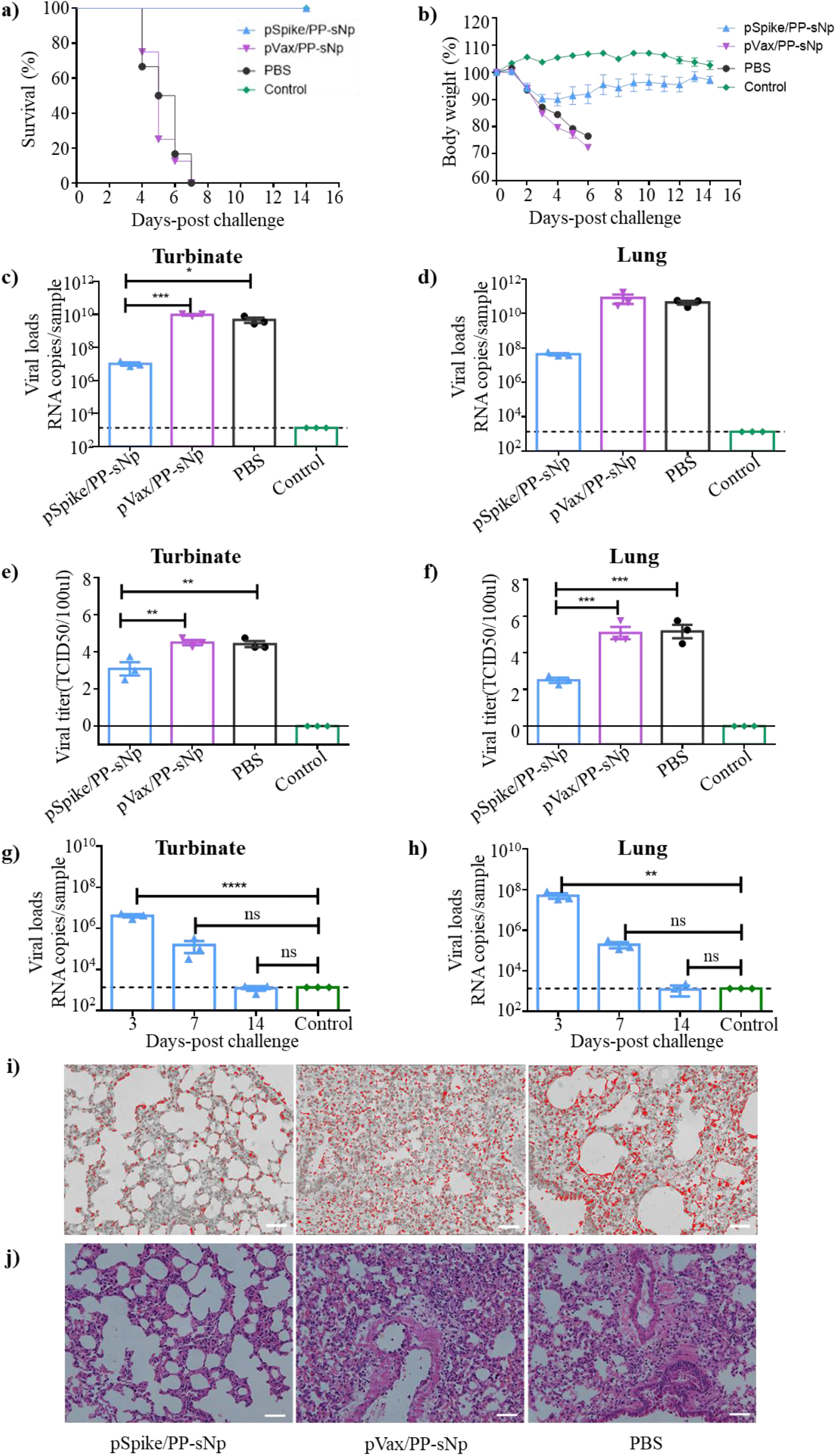
Pulmonary immunization with pSpike/PP-sNp confers complete protection against lethal SARS-CoV-2 challenge in mice. Forty days after the initial immunization, mice were challenged with a lethal dose (50 LD_50_) of SARS-CoV-2 C57MA14 strain via intranasal instillation, and the indicated tissues were collected at indicated time points after challenge to detect viral loads and the lung pathology. **a**) The survival rate of mice (n= 10 mice/group) intratracheally immunized with pSpike/PP-sNp, pVax/PP-sNp (Mock control) and PBS after the SARS-CoV-2 challenge. Untreated mice without challenge were served as a control (Control) **b**) The body weight changes of mice (n= 10 mice/group) intratracheally inoculated with pSpike/PP-sNp, pVax/PP-sNp and PBS after the SARS-CoV-2 challenging. Untreated mice without challenge were served as a control (Control, marked in green). Data represent mean ± SEM. Viral RNA loads in **c**) the turbinates and **d**) the lungs of mice treated by pSpike/PP-sNp, pVax/PP-sNp and PBS as described above and that in untreated counterpart (Control) 3 days after the SARS-CoV-2 challenge. Viral titers in **e**) the turbinates and **f**) the lungs of mice treated by indicated formulations as described above and that in untreated counterpart (Control) 3 days after the SARS-CoV-2 challenging. The viral loads in **g**) the turbinates and **h**) the lungs of mice treated by intratracheally vaccinated pSpike/PP-sNp 3, 7, 14 days post-challenge. **i**) Immunohistochemistry assay for SARS-CoV-2 N protein 3 days post-challenge. Scale bar, 100 μm. Positive signals are shown in red. **j**) Representative hematoxylin-eosin staining (H&E) staining of lung pathology 3 days after the SARS-CoV-2 challenge. Scale bar, 100 μm. Data in **c-h** represent mean ± SEM (n= 3 biologically independent samples). One-way ANOVA with Dunnett’s post-hoc test was used to determine significance (ns represent not significant, \**P*< 0.05, \*\**P*< 0.01, *\*\*\*P*< 0.001, *\*\*\*\*P*< 0.0001).

## Discussion

The development of mucosal COVID-19 vaccines that impart protective immunity would be highly desirable. However, apart from extensive investigations using adenovirus-based vaccines^17, 18, 41^, whether DNA vaccines could achieve this goal remains unclear. In this study, we demonstrate a COVID-19 DNA vaccine (pSpike/PP-sNp) delivered via pulmonary route could induce a robust protective immunity consisting of mucosal, humoral and cellular immune responses, which is sufficient to completely protect the upper and lower respiratory tracts against lethal SARS-CoV-2 challenge in mice. Early studies suggested that DNA vaccines are poorly immunogenic in the airways with low levels of antigen expression owing to the barriers posed by the respiratory tract. This could be reflected by the previous study indicating intranasal COVID-19 DNA vaccination were not able to elicit mucosal sIgA and circulating IgG against SARS-CoV-2 even after three times of repeated dosing^27^. Similar observations have been reported in another study showing that the serum neutralization IC_50_ of a COVID-19 DNA vaccine just reached 1/83.8 against SARS-CoV-2 pseudoviruses after seven-times repeated intranasal vaccinations^42^. Although the previous studies reveal the intranasal administered DNA vaccines could trigger a certain degree of SARS-CoV-2 specific immune responses^27, 42^, none of them has disclosed critical data regarding the mucosal application of COVID-19 DNA vaccine, including sIgA levels in BALF samples, details in the subsets of effector T cells and memory T cells, and the protection efficiency in a SARS-CoV-2 challenge assay.

In order to potentiate the mucosal COVID-19 DNA vaccine, we adopted the PP-sNp gene delivery system which is specifically developed for pulmonary DNA transfection^28, 32^. Previous studies from both our group and others have clearly demonstrated that poloxamine-based delivery system mediates significantly better levels of DNA transfection and lower levels of associated inflammatory response in the airways of mouse and pig models than were achieved with the cutting-edge lipid-based formulations^26, 32, 33^. PEI was adopted as a control formulation because it is one of the most well-studied polymer-based gene carrier and has been applied in many mucosal DNA vaccine applications^43^. Although PEI appears to be promising in delivery of mucosal DNA vaccines encoding antigens of Influenza, HIV and SARS-CoV^43^, we found it fails to mediate efficient DNA transfection in the mouse airway, perhaps owing to its poor mucus penetrating properties. We applied an improved method of preparing nanoparticles without aggregation to enhance the utility of PP-sNp for in vivo applications^44^. The optimized nanocomplexes appeared as evenly distributed spherical nanoparticles with sizes being smaller than mucus mesh pores (~140±50 nm)^45^. The unique high mobility in mucus gel facilitates deep penetration and widely spread of PP-sNp through the airway mucus layer (Fig. 6a), thereby improving the probability of DNA payloads encounter and uptake by target cells. Subsequently, the targeting moiety within PP-sNp (which binds the ICAM-1/CD54) provides the possibility to specifically deliver the DNA cargos via receptor-mediated uptake, concurrent with efficient endosome escape and nucleus localization in these cells^28^. By virtue of these merits, PP-sNp efficiently mediates the transfection of antigen-expressing plasmid in pulmonary DCs and AECs (Fig. 6b). Being the most powerful APCs, pulmonary DCs are imperative in shaping antigen-specific immune responses^46^. The enhanced DNA vaccine uptake by pulmonary DCs and subsequent DC differentiation may lead to more profound and durable antigen presentation to T-cells^47^. Our finding is also in agreement with previous evidences demonstrating a positive correlation between the ratio of antigen-activated DCs and the magnitude of T_EM_-biased response^48^. Meanwhile, AECs appear to be essential for determining the potency of mucosal immune response and IgA production. Several studies have suggested that AECs provides a constant supply of cytokines (such as IL-6) which are essential for B cell proliferation, IgA isotype switch and differentiation into IgA producing plasma cells^49, 50^. The close proximity of AECs to DCs and B cells situated in respiratory mucosa probably ensures efficient antigen cross-presentation which directs B cells isotype switch towards IgA1 and IgA2 with the help of cytokines produced by AECs^50^. AECs also hold critical roles in orchestrating innate and adaptive immune responses in the respiratory system during viral infection^51, 52^. As an RNA sequencing analysis reveals, pSpike/PP-sNp successfully activated innate and adaptive anti-virus pathways in vaccinated mice (Supplementary Fig. 22, more details are reported in the Supplementary Results), which is in consistent with previous findings suggesting the transiently activated innate immunity is sufficient to augment subsequent adapted immune responses^53^.

**Figure 6.**
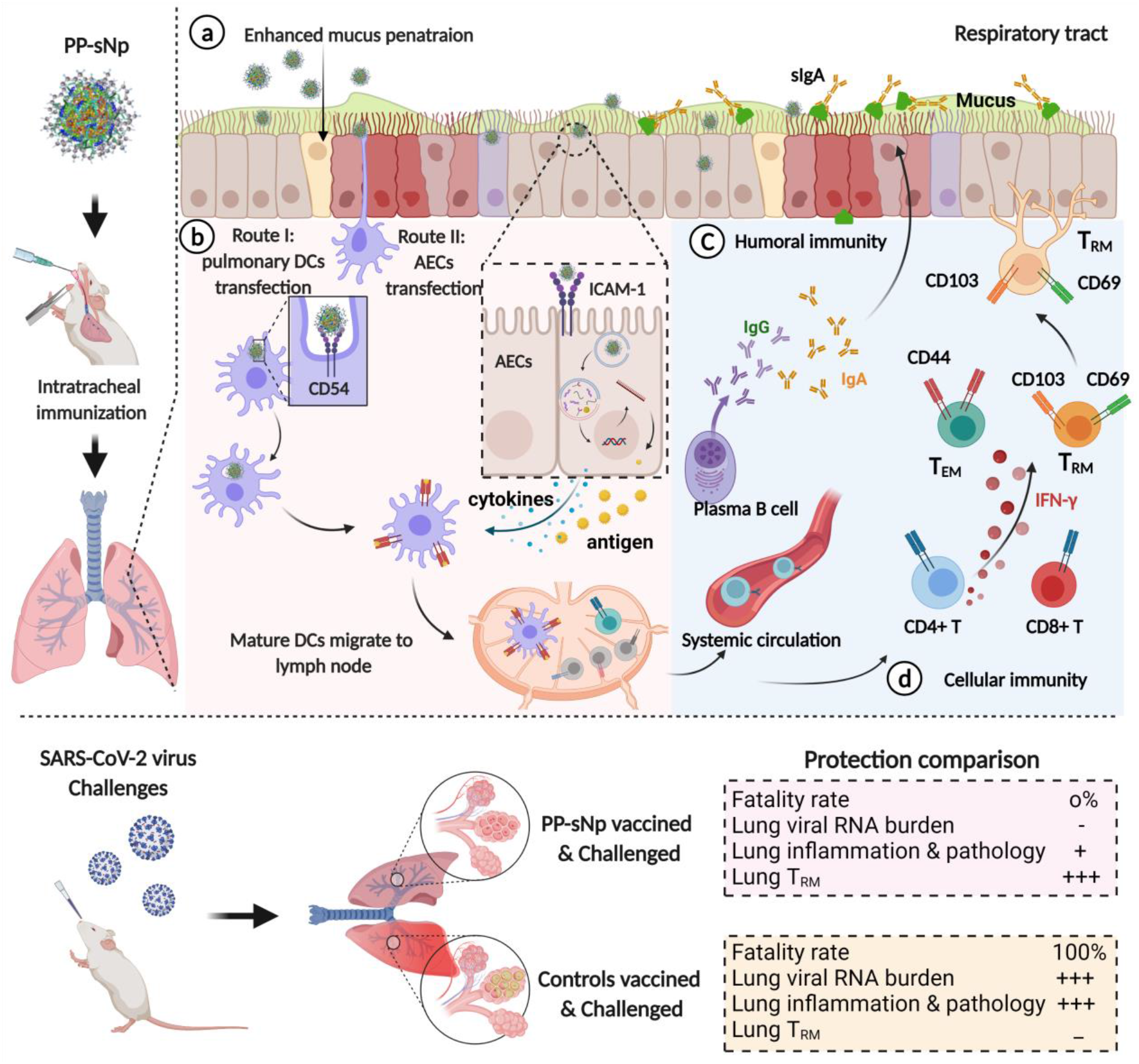
Schematic of the SARS-CoV-2 specific immune responses and subsequent protection established by pSpike/PP-sNp vaccine inoculated via respiratory route. **a**) After pulmonary inoculation, pSpike/PP-sNp is able to penetrate efficiently through the physical and biological barriers at the airway mucosal site. **b**) With the help of targeting moiety (ICAM-1/CD54 ligand) in PP-sNp, pSpike/PP-sNp is internalized and captured by airway epithelial cells (AECs) or pulmonary dendritic cells (DCs). The cationic moiety within PP-sNp mediates endosome escape and nucleus localization of pSpike, followed by the transcription and translation process. The SARS-CoV-2 derived antigens expressed in AECs are subsequently presented to other immune cells such as pulmonary DCs. AECs simultaneously orchestrate the adaptive immune responses via the transient secretion of cytokines. Mature pulmonary DCs then migrate to the bronchial-associated lymphoid tissues (e.g., mediastinal lymph node) and present antigens to naïve T cells and B cells for humoral and cellular immune responses. **c**) Activated B cells proliferate and differentiate into antibody-secreting plasma cells to generate secretory IgA (sIgA) and systemic IgG antibodies, the former of which efficiently neutralizes invading SARS-CoV-2 within the upper and lower respiratory tracts. **d**) Meanwhile, a portion of T cells obtains a tissue resident memory phenotype (TRM), enabling them to reside in the airway and respond rapidly when encountering SARS-CoV-2 virus. Other activated CD4^+^ or CD8^+^ T cells, including CD44^hi^CD62L^lo^ T cells, CD4^+^IFN-γ T cells and CD8^+^IFN-γ T cells, take part in the process of eliminating SARS-CoV-2 infections as well. The robust and comprehensive immunity conferred by pulmonary vaccination of pSpike/PP-sNp probably controls SARS-CoV-2 replication and removes the viruses at the initial sites of infection, thus protects the vaccinated mice from lung lesions and death.

The complete protection against lethal SARS-CoV-2 challenge was largely due to the ability of pSpike/PP-sNp in stimulating comprehensive mucosal, humoral and cellular immunity, characterized by robust secretion of sIgA and NAb as well as potent T cell responses (especially T_RM_ cells) in the respiratory system. pSpike/PP-sNp efficiently activated systemic immune response, concurrent with significant levels (ND_50_ at ~1/1875) of NAb in serum. Previous findings suggest a serum NAb titer >300 was generally associated with protection against SARS-CoV-2^54^. It is worth noting that pSpike/PP-sNp vaccinated mice displayed B cells secreting IgA and high levels of SARS-CoV-2-specific sIgA antibody titer (with a mean dilution titer of 1/102.4) in the BALF samples (Fig. 6c), which are even higher than those induced by intranasal vaccination of adjuvanted subunit vaccines according to previous publications^55–57^. Although the SARS-CoV-2-specific sIgA antibody titer in the BALF of intranasally vaccinated mice using adenovirus-vector is insurmountable when compared to non-viral vaccine induced counterpart, substantial NAb titers could be detected in BALF samples of pSpike/PP-sNp vaccinated mice with a level (~1/73) that is comparable to the adenovirus-based counterpart (~1/100)^19^.

Additionally, it has been long recognized that high-quality antigen-specific T cell responses are pivotal in combating various types of coronavirus infection^9^. pSpike/PP-sNp successfully mediated the presence of SARS-CoV-2 specific CD8^+^IFN-γ T cells (cytotoxic T cells) and elicited a Th1-biased cellular immune responses (Fig. 6d), both of which would be advantageous in eliminating coronavirus without adverse effects, since Th2 cell responses are suggested to be associated with enhancement of lung diseases^58^. pSpike/PP-sNp also generated a robust memory-based cellular immunity in the airways, including the significantly activated T_EM_ cells and T_RM_ cells with a resident memory phenotype (Fig. 6d). T_EM_-biased response created by pSpike/PP-sNp, which could immediately act on and rapidly removes respiratory pathogens, is particularly pronounced in the lung. But we failed to observe increased ratio of T_CM_ cells, which is in line with previous findings indicating that T_CM_-biased response generally induced by conventional route of vaccination (e.g., electroporation)^59^. Accumulating evidences demonstrate that pulmonary CD4^+^T_RM_ and CD8^+^T_RM_ cells play crucial roles in resisting respiratory pathogen infections, including MERS-CoV, SARS-CoV and SARS-CoV-2^3, 60^. The airway T_RM_ cells, which are exclusively generated through the respiratory route vaccination/infection, reside in the lung and does not recirculate^61^, so that they can instantly recognize invading pathogens in the airway and efficiently prevent virus replication at the early stage of infection/re-infection.

Despite the promising results we observed in the current study, pSpike/PP-sNp is still far from successful clinical translation and more in-depth investigations are necessary. There are several limitations that we did not address in this study and will be useful topics for future studies, including the absence of data on the neutralization and protection efficiency elicited by pSpike/PP-sNp against emerging SARS-CoV-2 variants of concern. Similar to those cases of authorized COVID-19 vaccines^62^, the neutralizing activity of NAb induced by the pSpike/PP-sNp vaccine may suffer a significant decrease within several months/years after vaccination, more boost doses may be necessary. Besides, immunization and challenge studies with larger animals such as non-human primates should be carried out to confirm the extent of protective mucosal immunity conferred by pSpike/PP-sNp. Another limitation relates to the intratracheal dosing which is not appropriate to be applied in humans when considering its poor compliance. Most of the relevant studies chose the intranasal inoculation because of its noninvasive and convenient features, but there are still huge concerns and uncertainties regarding intranasal route of vaccination. For example, negative perception for nasal vaccines was generated from reported cases of Bell’s palsy after intranasal dosing of influenza vaccines^63, 64^. Alternatively, the noninvasive nebulized formulations seem to be one of the most appropriate approaches in delivering mucosal vaccines to the human airway. However, the nebulized DNA formulations still face many challenges as indicated by a previous study showing that as little as 10% of the DNA payload in a nebulization device chamber could be successfully emitted^65^, it thus requires advanced nebulization strategies and further optimization on the formulations to ensure the transfection efficiency^66^. Finally, the mechanism and pathways that involved in the pSpike/PP-sNp induced immune responses will be explored in details with the help of cutting-edge technologies such as the single-cell RNA sequencing in order to further improve the vector/platform design and our understanding of DNA-based mucosal vaccines.

In summary, our dataset reveals that pSpike/PP-sNp with pulmonary mucus penetrating properties was capable of inducing comprehensive mucosal, humoral and cellular immune responses to provide complete protection against SARS-CoV-2 infection. The safe profile and ability to potentiate DNA vaccines for strong mucosal immunity make PP-sNp a promising platform for COVID-19 mucosal vaccines if its efficacy can be shown in large animal models and clinical trials, since robust protection at the respiratory mucosal site would be, at least theoretically, one of the most effective means to prevent the infection of SARS-CoV-2.

## ACKNOWLEDGMENTS

This work was supported by the National Natural Science Foundation of China (NSFC, Grant No. 82041045, 82173764 and 31972720), the major project of Study on Pathogenesis and Epidemic Prevention Technology System (2021YFC2302500) by the Ministry of Science and Technology of China, the Chongqing Talents: Exceptional Young Talents Project (CQYC202005027) and the Natural Science Foundation of Chongqing (cstc2021jcyj-msxmX0136). We also thank H. Zeng, Y. Zhuang, J. Gu, Jinyong Zhang, L. Peng, H. Sun and Jianxiang Zhang (Third Military Medical University, Chongqing, China) for helpful discussions and careful proofreading of the manuscript.

## AUTHOR CONTRIBUTIONS

S.G. and Q.Z. conceived and directed the project. Y.D., D.L., W.Z., P.L., P.C., B.P. and J.R. contributed experimental materials. S.S., J.T., Q.Z., L.C., C.X., C.S., Y.O., C.L., H.L., and Y.D. performed experiments and analyzed data. E.L., Y.G., C.T. and Y.L. designed and performed the challenge study and relevant end-point investigations. C.L. was responsible for the bioinformatics and RNA sequencing data analysis. S.G., G.Z., B.W., Y.L. and Q.Z. designed and supervised the research. S.G. and S.S. wrote the manuscripts with help and comments from all authors.

## COMPETING FINANCIAL INTERESTS

S.G., S.S., Q.Z. and C.L. have applied for patents related to this study. B.W. is a scientific co-founder of the biotechnology company Advaccine Biopharmaceuticals (Suzhou, China), which focuses on the development of DNA vaccines. G.Z., C.S. and Y.D. are employees of Advaccine Biopharmaceuticals Co., Ltd. B.P. is a scientific co-founder of In-Cell-Art (Nantes, France) and owns stock of In-Cell-Art, which commercializes tetra-functional block copolymers for DNA vaccines. The other authors declare that they have no competing financial interests.

## Methods

### Reagents

Poloxamine 704 (T704) was kindly provided by InCellArt (Nantes, France). All synthetic peptides were manually synthesized by Chinese Peptide Company (Hangzhou, China) with purity > 95%. Branched PEI (average molecular weight at 25 kDa), Cell Counting Kit-8 (CA1210) and D-Luciferin (L6882) were all purchased from Sigma-Aldrich (Saint Louis, MO, USA). Lipofectamine2000 was purchased from Invitrogen (11668019, Carlsbad, USA). pGL4.51-Luciferase Reporter Vectors (pFLuc, E1320) was purchased from Promega (Madison, USA). Plasmids encoding SARS-CoV-2 S protein (pSpike) and pVax were kindly provided by Advaccine Biopharmaceuticals Co., Ltd (Suzhou, China). Purified full length S1 + S2 ECD spike protein of SARS-CoV-2 was purchased from Sino Biologics (40589-v08B1, Beijing, China). For in vivo uptake and flow cytometric analysis, plasmid was fluorescently labeled with the Cy5 fluorophores using the Mirus Label IT® tracker intracellular nucleic acid localization kit (MIR 7021, Mirus Bio, Madison, USA) according to the manufacturer’s instruction. Other reagents were obtained from Sigma-Aldrich (Saint Louis, MO, USA) as analytical grade or better.

### Cell lines

MH-S cell line (Mice alveolar macrophages cells), DC2.4 cell line (Mouse bone marrow-derived dendritic cells), BEAS-2B (human bronchial epithelial cells) cell line and Calu-3 (human lung cancer cells) cell line were obtained from ATCC (Manassas, VA, USA). 16HBE (human bronchial epithelial cells) cell line was generously provided by Prof. Dr. Dieter C. Gruenert (University of California at San Francisco, CA, USA). ACE2-293T cells (ACE2-expressing cell line, constructed by hygromycin B screening) were purchased from PackGene (LV-2058, Guangzhou, China). Cells were maintained in medium DMEM (Gibco, USA) supplemented with 10% fetal bovine serum (Gibco, USA), penicillin (100 units/mL) and streptomycin (100 μg/mL) (complete medium) at 37 °C in 5% CO2. All cell lines used in current study were obtained from original providers who authenticated the cell lines using morphology, karyotyping and PCR-based approaches. No additional authentication has been performed. All cell lines tested negative for mycoplasma contamination. All experiments were performed on cells in the logarithmic growth phase.

### Preparation and characterization of pDNA/PP-sNp formulations

The pDNA/PP-sNp formulations were prepared via a simple self-assembly process as described previously^28^. For in vivo applications, T704 stock solution (10 mg/mL) prepared in nuclease-free water was mixed with an equal volume of peptide solution (0.667 mg/mL) using a self-designed microfluidic mixer. pDNA solution (0.6 mg/mL) in double volumes were mixed with the above T704/peptide solution using the same mixer. The complex was incubated for 20 min at room temperature before further use. For in vitro study, T704 solution at a certain concentration (defined as the w/w ratio between T704 and pDNA) and peptide solution at a specific concentration (defined as the N/P ratio; namely, the ratio between nitrogen residues in peptide and nucleic acid phosphate groups) were applied using similar methods. For PEI based formulation, brPEI (25 kDa, Sigma) at an optimum N/P ratio of 10 was mixed with pDNA solution in nuclease-free water, the resulting complex was incubated at room temperature for 20 min and served as a polyplex control. the Lipofectamine 2000 complex, serving as a lipoplex control, was prepared according to the manufacturer’s instructions using the optimum concentration. Size measurements were performed using dynamic light scattering (DLS) on a Malvern Zetasizer Nano-ZS (Malvern). The morphology of pDNA/PP-sNp formulations were investigated by transmission electron microscope and scanning electron microscope (JEOL Ltd).

### Cellular uptake and in vitro transfection

For cellular uptake investigation, the Cy5 conjugated pDNA was prepared according to manual instructions. Cells were incubated with Cy5-pDNA/PP-sNp or naked Cy5-pDNA for 4 h in Opti-MEM I Reduced Serum Medium (31985062, Invitrogen). The cells were further collected for the analysis of mean fluorescence intensity (MFI) by flow cytometry. To evaluate the in vivo transfection of PP-sNp, pDNA/PP-sNp complexes encoding firefly luciferase (i.e., pFLuc/PP-sNp) were prepared. pFLuc/PP-sNp and naked-pFLuc were incubated with cells for 4 h in Opti-MEM I Reduced Serum Medium, then were replaced by fresh complete medium. The protein expression of firefly luciferase in cells was observed by bioluminescence imaging at 48 h by Firefly Luciferase Reporter Gene Assay Kit (RG005, Beyotime).

### Optical video recording of PP-sNp and the mean-squared displacement (MSD) analysis

Leica SP8 microscope equipped with a 40× water objective was used to record the motion of PP-sNp. Cy5-DNA-containing PP-sNp were prepared as described above. PP-sNp (20 μL, 1.3 μg/μL) were mixed with mucin gel (400 μL, Mucin II solution 3%, *w/v*) mimicking mucus, followed by a 30 min incubation at 37°C. DNA/PEI nanoparticles were also added as control. Fifteen-second Movies (Frame rate 23 fps) were captured using the LAS4.5 software (Leica). The trajectories of the nanoparticles were precisely quantified from the videos by software (TrackMate plugin in FIJI (ImageJ)), then the trajectory data was used to calculate the MSD and the corresponding diffusion coefficients (*De*) in MATLAB through the following equations, as implemented in MSD Analyzer.

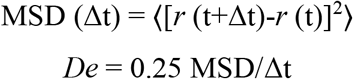

where *De* represents the effective diffusion coefficient and Δt represents the time interval.

### Animals

6–8 weeks old specific pathogen-free (SPF) female BALB/c mice were purchased from the Beijing HFK Bioscience Co., Ltd (Beijing, China). All animal studies were approved by the Laboratory Animal Welfare and Ethics Committee of Third Military Medical University and were performed in accordance with the institutional and national policies and guidelines for the use of laboratory animals. The mice were kept and vaccinated in SPF facilities, and provided with free access to sterile food and water. Animals were randomly divided into groups and conceded an adaption time of at least 7 days before the beginning of the experiments. For intratracheal dosing, the mice were sedated with isoflurane and received the formulations via a self-improved micro-injector.

### Bioluminescence imaging

Mice were anaesthetized with isoflurane, followed by an injection of the substrate D-Luciferin (150 mg/kg) intraperitoneally. Bioluminescence was measured 10 min later using a Lumina Series III In Vivo Imaging System (PerkinElmer). To evaluated the in vivo delivery capability and visualize protein expression and tissue distribution of pDNA/PP-sNp formulations, pDNA encoding a firefly luciferase (pFLuc) was incorporated. The formulations were administered to mice via intratracheal (i.t.) route. The protein expression of firefly luciferase mediated by pFLuc/PP-sNp was observed by bioluminescence imaging at 24 h, 48 h, 72 h and 7 days post-dosing, respectively. The pFLuc/PEI (at N/P ratios of 10) and naked pFLuc plasmid counterparts were also injected via i.t. route with same dose of pFLuc to evaluate the bioluminescence in vivo.

### Bone marrow derived dendritic cells (BMDC) maturation study

Bone marrow cells were isolated from the femurs of female BALB/c mice and cultured in RPMI 1640 complete medium (Gibco, USA) supplemented with 10% FBS, 1% penicillin/streptomycin, 10 ng/mL of Interleukin-4 (IL-4) and Granulocyte-Macrophage Colony Stimulating Factor (GM-CSF). The culture media was replaced with fresh media on day 2 and 5 to remove the non-adherent and loosely adherent cells. The remaining cells continued to culture for another 2 days. To examine the maturation of BMDCs in vitro, BMDCs (1 × 10^6^ mL^-1^) were co-cultured with pSpike/PP-sNp and naked-pSpike only for 24 h, respectively. Subsequently, FITC anti-mouse CD11c (117305, Biolegend), PE-Cy7 anti-mouse MHC-II (25-5321-82, Invitrogen), and APC anti-mouse CD86 (105011, Biolegend) were used to stain the cells in flow cytometry staining (FACS) buffer for 30 min at 4 °C before being washed and analyzed by BD FACS Array software™ on a BD FACS Array flow cytometer (BD Biosciences, USA).

### Mouse vaccination and challenge experiments

Mice were immunized on day 0 and boosted with the same dose on day 14 and 28, respectively. Each anesthetized mouse intratracheally received 50 μL of pSpike/PP-sNp formulation containing 15 μg pSpike. pVax/PP-sNp and phosphate buffered saline (PBS) was adopted as a mock control and a negative control, respectively. Mice were sacrificed on day 35 for assessing respiratory mucosal immune response, cellular immune response and memory establishment. Relevant tissues (lung and spleen) were harvested and processed for flow cytometry analysis. Bronchoalveolar lavage fluid (BALF) was collected by washing the lungs of euthanized mice with 500 μL of ice-cold PBS containing 0.05% Tween-20. Collected BALF was stored at −80°C for further use.

The SARS-CoV-2 challenge model was based on a novel mouse-adapted SARS-CoV-2 strain, C57MA14 (NCBI GenBank number: OL913104.1, details can be found in: https://www.ncbi.nlm.nih.gov/nuccore/2167992552), that causes severe respiratory symptoms, and mortality to BALB/c mice. Immunized BALB/c mice were challenged intranasally with 50 LD_50_ SARS-CoV-2 C57MA14 on day 40 post initial immunization. On day 3 post challenge, 3 animals/group were sacrificed, and the lung and trachea tissues were collected for subsequent viral loads detection.

### Quantification of viral by quantitative RT-PCR and TCID50 in challenged mouse tissues

Viral RNA in lung and turbinate tissues from challenged mice was detected by quantitative reverse transcription PCR (RT-qPCR). Briefly, tissue samples were homogenized with stainless steel beads in a Tissuelyser-24 (Shanghai jingxin Industrial Development CO., LTD) in 500 uL of DMEM. Viral RNA in tissues were divided into two parts. One part was extracted using the QIAamp Viral RNA Mini Kit (QIAGEN) according to the manufacturer’s protocol. SARS-CoV-2 RNA quantification was performed by RT-qPCR targeting the S gene of SARS-CoV-2 using One Step PrimeScript RT-PCR Kit (Takara) with the following SARS-CoV-2 specific primers and probes: CoV-F3 (5’-TCCTGGTGATTCTTCTTCAGGT-30), CoV-R3 (5’-TCTGAGAGAGGGTCAAGTGC-30), and CoV-P3 (5’-FAM-AGCTGCAGCAC CAGCTGTCCA-BHQ1-30). Another part was serially diluted in DMEM and added into Vero E6 cells in 96-well plates. The plates were incubated 1 hour at 37°C with 5% CO_2_, the inoculation was replaced with DMEM containing 2% FBS and 1% penicillin-streptomycin. After incubating for 72 h, the median tissue culture infective dose (TCID50) was detected by the cytopathic effect (CPE).

### Enzyme linked immunosorbent assay (ELISA)

SARS-CoV-2 S protein specific antibodies in mouse serum and BALF were measured by ELISA. Briefly, polystyrene microtiter 96-well plates were coated with full length SARS-CoV-2 spike protein (3μg/mL in carbonate buffer, pH=9.6) and incubated overnight at 4 °C. After blocking with 1% bovine albumin (BSA) in PBS, 100 μl/well pre-diluted samples were added into the plates with 1 h incubation at 37 °C. After three-times washes with PBST (PBS with 0.05% Tween-20), plates were added with horseradish peroxidase (HRP) conjugated goat anti-mouse IgG (Ab231712), IgA (Ab97235), IgG1 (ab97240), or IgG2a (Ab97245) (1:10000, Abcam, UK) and incubated for 40 min at 37 °C. Plates were then washed three-times and added with peroxidase substrate (Ab171522, Abcam, UK), the reaction was terminated by stop solution (Ab171529, Abcam, UK) and the absorbance at 450 nm was read using a microplate reader (AID iSpot, Germany).

### SARS-CoV-2 pseudovirus neutralization assay

Mouse serum samples from pSpike/PP-sNp vaccinated mice were serially diluted double fold starting at a 1:100 dilution with DMEM contain 2% FBS for the assay. Serum or BALF were incubated with 10 μl of Luc-SARS-Cov-2 pseudotyped virus (LV-2058, PackGene, China) for 60 min, then added to the HEK293T cells stably expressing ACE2 to incubate in a standard incubator (37°C, 5% CO_2_) for 72 h. Post infection, cells were lysed and detected using a luminescence reporter gene assay system (RG006, Beyotime Biotechnology, China). The Luciferase activity was measured using the Promega GloMax Navigator Detection System (GloMax, Promega, USA) and expressed as relative light units (RLU). The 50% neutralization titers (NT_50_) were calculated as the serum dilution at which RLU were reduced by 50% compared with RLU in virus control wells after subtraction of background RLU in cell control wells.

### Tissue processing and flow cytometry

Single cell suspensions of splenic and pulmonary lymphocytes were prepared from resected spleens and lungs of mice. The lung tissues were cut into scraps and digested with collagenase ⊓ (0.5 mg/mL, C8150, Solarbio, China) in calcium chloride (1mM) and magnesium chloride (1mM) solution at 37 °C for 1 h with shaking at 220 rpm. Next, the samples were filtered through a 75 mm cell strainer to obtain a single cell suspension. The immune cells were obtained by density gradient centrifugation with Percoll (17-0891-09, GE Healthcare, USA) according to the manufacture’s instruction. Splenic lymphocytes were collected by grinding spleen in PBS then passed through a 75 mm cell strainer, cell pellets were re-suspended in 5 mL of red blood cell lysis buffer (RT122-02, TIANGEN, China) for 5 min at RT for remove the red blood cell. PBS was added to wash the cells twice, then centrifuged at 1500 × g for 5 min, the cell pellets were eventually re-suspended in RPMI1640 media supplemented with 10% FBS and 1% penicillin/streptomycin.

For flow cytometric study, cells were first stained with the LIVE/DEAD fixable cell stains kit (65-0865-14, Invitrogen, USA) according to the manufacturer’s protocol. For surface markers, the cells were incubated with anti-mouse CD4 (11-0041-82, Invitrogen), anti-mouse CD8a (45-0081-82, Invitrogen), For intracellular cytokine staining, cells were stimulated with the overlapping peptide pool spanning of 14-mer peptides overlapping by nine amino acids from the SARS-CoV-2 RBD proteins (see Supplementary Notes) for 6 h at 37 °C, 5% CO_2_. Then the cells were incubated with anti-mouse IL-4 (17-7041-82, Invitrogen) and anti-mouse IFN-γ (12-7311-82, Invitrogen) after processing with the Cytofix/Cytoperm Fixation/Permeabilization Kit (554714, BD Biosciences,) according to the manufacturer’s instructions. In order to detect the T_EM_/T_CM_ and T_RM_ cells, the cell samples were stained with the following indicated antibodies in FACS buffer: anti-CD62L (161204, BioLegend), anti-CD44 (25-0441-82, BioLegend), anti-CD69 (104506, BioLegend) and anti-CD103 (121416, BioLegend). All the samples were measured on a BD FACS Array flow cytometer (BD Biosciences). Data are analyzed with FlowJo software V10.

### Enzyme linked immunospot (ELISpot) assay

Cellular immune responses in mice were performed using mouse IFN-γ/IL-4 ELISPOT PLUS plates (3321-4AST-2/3311-4APW-2, MABTECH, Sweden). 96-well ELISPOT plates were pre-treated as the manufacturer’s instructions. 5 ×10^5^ mouse splenocytes or pulmonary lymphocytes were plated into each well and stimulated with the above-mentioned peptide pools at a final concentration of 1 μg of each peptide per well. Additionally, PMA/Ionomycin were added as a positive control and RPMI 1640 media was used as a negative control. After incubation at 37 °C, 5% CO_2_ for 24 h, the plates were washed with PBS and incubation with biotinylated anti-mouse IFN-γ or IL-4 antibody for 2 h at RT. Finally, TMB substrate solution were added to visualize the spots. Spots were scanned and quantified by an ImmunoSpot CTL (Bio-Rad) reader. Spot-forming unit (SFU) per million cells was calculated by subtracting the negative control wells.

### Statistics and analysis

Statistical analyses were performed using the GraphPad Prism 8 (GraphPad Software, USA). Unless otherwise specified, data were expressed as mean ± Standard Error of Mean (SEM). Dual comparisons were made using Welch’s t-test, and comparisons between multiple conditions were analyzed using analysis of variance (ANOVA) followed by the appropriate post-hoc tests. All the tests were two tailed. Differences were considered statistically significant when *P* < 0.05. All of the experiments were successfully repeated at least twice with three or more biological replicates to ensure the reproducibility of the data.

## References

1. Azzi, L. et al. Mucosal immune response in BNT162b2 COVID-19 vaccine recipients. EBioMedicine 75, 103788 (2022).

2. Zhou, D. et al. Robust SARS-CoV-2 infection in nasal turbinates after treatment with systemic neutralizing antibodies. Cell Host Microbe 29, 551–563 e555 (2021).

3. Afkhami, S. et al. Respiratory mucosal delivery of next-generation COVID-19 vaccine provides robust protection against both ancestral and variant strains of SARS-CoV-2. Cell 185, 896–915 e819 (2022).

4. Bricker, T.L. et al. A single intranasal or intramuscular immunization with chimpanzee adenovirus-vectored SARS-CoV-2 vaccine protects against pneumonia in hamsters. Cell Rep 36, 109400 (2021).

5. Hassan, A.O. et al. An intranasal vaccine durably protects against SARS-CoV-2 variants in mice. Cell Rep 36, 109452 (2021).

6. King, R.G. et al. Single-Dose Intranasal Administration of AdCOVID Elicits Systemic and Mucosal Immunity against SARS-CoV-2 and Fully Protects Mice from Lethal Challenge. Vaccines (Basel) 9 (2021).

7. Li, Y., Jin, L. & Chen, T. The Effects of Secretory IgA in the Mucosal Immune System. Biomed Res Int 2020, 2032057 (2020).

8. Braun, J. et al. SARS-CoV-2-reactive T cells in healthy donors and patients with COVID-19. Nature 587, 270–274 (2020).

9. Sekine, T. et al. Robust T Cell Immunity in Convalescent Individuals with Asymptomatic or Mild COVID-19. Cell 183, 158–168 e114 (2020).

10. Wang, Z. et al. Enhanced SARS-CoV-2 neutralization by dimeric IgA. Science Translational Medicine 13, eabf1555 (2021).

11. Sterlin, D. et al. IgA dominates the early neutralizing antibody response to SARS-CoV-2. Science Translational Medicine 13, eabd2223 (2021).

12. Grau-Exposito, J. et al. Peripheral and lung resident memory T cell responses against SARS-CoV-2. Nat Commun 12, 3010 (2021).

13. Shakya, A.K., Chowdhury, M.Y.E., Tao, W. & Gill, H.S. Mucosal vaccine delivery: Current state and a pediatric perspective. J Control Release 240, 394–413 (2016).

14. Mudgal, R., Nehul, S. & Tomar, S. Prospects for mucosal vaccine: shutting the door on SARS-CoV-2. Hum Vaccin Immunother 16, 2921–2931 (2020).

15. Miquel-Clopes, A., Bentley, E.G., Stewart, J.P. & Carding, S.R. Mucosal vaccines and technology. Clin Exp Immunol 196, 205–214 (2019).

16. Lavelle, E.C. & Ward, R.W. Mucosal vaccines - fortifying the frontiers. Nat Rev Immunol 22, 236–250 (2022).

17. Wu, S. et al. A single dose of an adenovirus-vectored vaccine provides protection against SARS-CoV-2 challenge. Nat Commun 11, 4081 (2020).

18. Doremalen, N.v. et al. Intranasal ChAdOx1 nCoV-19/AZD1222 vaccination reduces viral shedding after SARS-CoV-2 D614G challenge in preclinical models. Science Translational Medicine 13, eabh0755 (2021).

19. Hassan, A.O. et al. A Single-Dose Intranasal ChAd Vaccine Protects Upper and Lower Respiratory Tracts against SARS-CoV-2. Cell 183, 169–184.e113 (2020).

20. See, I. et al. US Case Reports of Cerebral Venous Sinus Thrombosis With Thrombocytopenia After Ad26.COV2.S Vaccination, March 2 to April 21, 2021. JAMA 325, 2448–2456 (2021).

21. Park, K.S., Sun, X., Aikins, M.E. & Moon, J.J. Non-viral COVID-19 vaccine delivery systems. Adv Drug Deliv Rev 169, 137–151 (2021).

22. Tebas, P. et al. Safety and immunogenicity of INO-4800 DNA vaccine against SARS-CoV-2: A preliminary report of an open-label, Phase 1 clinical trial. EClinicalMedicine 31, 100689 (2021).

23. Modjarrad, K. et al. Safety and immunogenicity of an anti-Middle East respiratory syndrome coronavirus DNA vaccine: a phase 1, open-label, single-arm, dose-escalation trial. The Lancet Infectious Diseases 19, 1013–1022 (2019).

24. Silveira, M.M., Moreira, G. & Mendonca, M. DNA vaccines against COVID-19: Perspectives and challenges. Life Sci 267, 118919 (2021).

25. Suk, J.S. et al. Lung gene therapy with highly compacted DNA nanoparticles that overcome the mucus barrier. J Control Release 178, 8–17 (2014).

26. Alton, E.W.F.W. et al. Repeated nebulisation of non-viral CFTR gene therapy in patients with cystic fibrosis: a randomised, double-blind, placebo-controlled, phase 2b trial. The Lancet Respiratory Medicine 3, 684–691 (2015).

27. Chandrasekar, S.S. et al. Localized and Systemic Immune Responses against SARS-CoV-2 Following Mucosal Immunization. Vaccines (Basel) 9 (2021).

28. Guan, S. et al. Self-assembled peptide–poloxamine nanoparticles enable in vitro and in vivo genome restoration for cystic fibrosis. Nature Nanotechnology 14, 287–297 (2019).

29. Bui, T.M., Wiesolek, H.L. & Sumagin, R. ICAM-1: A master regulator of cellular responses in inflammation, injury resolution, and tumorigenesis. J Leukoc Biol 108, 787–799 (2020).

30. Iwasaki, A., Foxman, E.F. & Molony, R.D. Early local immune defences in the respiratory tract. Nat Rev Immunol 17, 7–20 (2017).

31. Braciale, T.J., Sun, J. & Kim, T.S. Regulating the adaptive immune response to respiratory virus infection. Nat Rev Immunol 12, 295–305 (2012).

32. Richard-Fiardo, P. et al. Evaluation of tetrafunctional block copolymers as synthetic vectors for lung gene transfer. Biomaterials 45, 10–17 (2015).

33. Caballero, I. et al. Tetrafunctional Block Copolymers Promote Lung Gene Transfer in Newborn Piglets. Molecular Therapy - Nucleic Acids 16, 186–193 (2019).

34. Shim, B.-S. et al. Intranasal immunization with plasmid DNA encoding spike protein of SARS-coronavirus/polyethylenimine nanoparticles elicits antigen-specific humoral and cellular immune responses. BMC Immunology 11, 65 (2010).

35. Torrieri-Dramard, L. et al. Intranasal DNA Vaccination Induces Potent Mucosal and Systemic Immune Responses and Cross-protective Immunity Against Influenza Viruses. Molecular Therapy 19, 602–611 (2011).

36. Bivas-Benita, M. et al. Airway CD8+ T cells induced by pulmonary DNA immunization mediate protective anti-viral immunity. Mucosal Immunology 6, 156–166 (2013).

37. Regnstrom, K. et al. PEI - a potent, but not harmless, mucosal immuno-stimulator of mixed T-helper cell response and FasL-mediated cell death in mice. Gene Ther 10, 1575–1583 (2003).

38. Blank, F. et al. Size-Dependent Uptake of Particles by Pulmonary Antigen-Presenting Cell Populations and Trafficking to Regional Lymph Nodes. American Journal of Respiratory Cell and Molecular Biology 49, 67–77 (2013).

39. Wilk, M.M. et al. Lung CD4 Tissue-Resident Memory T Cells Mediate Adaptive Immunity Induced by Previous Infection of Mice with Bordetella pertussis. J Immunol 199, 233–243 (2017).

40. Busch, D.H., Frassle, S.P., Sommermeyer, D., Buchholz, V.R. & Riddell, S.R. Role of memory T cell subsets for adoptive immunotherapy. Semin Immunol 28, 28–34 (2016).

41. Wu, S. et al. Safety, tolerability, and immunogenicity of an aerosolised adenovirus type-5 vector-based COVID-19 vaccine (Ad5-nCoV) in adults: preliminary report of an open-label and randomised phase 1 clinical trial. The Lancet Infectious Diseases 21, 1654–1664 (2021).

42. Kumar, U.S., Afjei, R., Ferrara, K., Massoud, T.F. & Paulmurugan, R. Gold-Nanostar-Chitosan-Mediated Delivery of SARS-CoV-2 DNA Vaccine for Respiratory Mucosal Immunization: Development and Proof-of-Principle. ACS Nano (2021).

43. Tang, J. et al. Nanotechnologies in Delivery of DNA and mRNA Vaccines to the Nasal and Pulmonary Mucosa. Nanomaterials (Basel) 12 (2022).

44. Vauthier, C., Cabane, B. & Labarre, D. How to concentrate nanoparticles and avoid aggregation? Eur J Pharm Biopharm 69, 466–475 (2008).

45. Suk, J.S. et al. The penetration of fresh undiluted sputum expectorated by cystic fibrosis patients by non-adhesive polymer nanoparticles. Biomaterials 30, 2591–2597 (2009).

46. Nguyen, T.L., Yin, Y., Choi, Y., Jeong, J.H. & Kim, J. Enhanced Cancer DNA Vaccine via Direct Transfection to Host Dendritic Cells Recruited in Injectable Scaffolds. ACS Nano 14, 11623–11636 (2020).

47. Kim, Y.C. et al. Strategy to Enhance Dendritic Cell-Mediated DNA Vaccination in the Lung. Advanced Therapeutics 3, 2000013 (2020).

48. Marzo, A.L. et al. Initial T cell frequency dictates memory CD8+ T cell lineage commitment. Nat Immunol 6, 793–799 (2005).

49. Beagley, K.W. et al. Interleukins and IgA synthesis. Human and murine interleukin 6 induce high rate IgA secretion in IgA-committed B cells. Journal of Experimental Medicine 169, 2133–2148 (1989).

50. Salvi, S. & Holgate, S.T. Could the airway epithelium play an important role in mucosal immunoglobulin A production? Clinical & Experimental Allergy 29, 1597–1605 (1999).

51. Whitsett, J.A. & Alenghat, T. Respiratory epithelial cells orchestrate pulmonary innate immunity. Nature Immunology 16, 27–35 (2014).

52. Saenz, S.A., Taylor, B.C. & Artis, D. Welcome to the neighborhood: epithelial cell-derived cytokines license innate and adaptive immune responses at mucosal sites. Immunological Reviews 226, 172–190 (2008).

53. Wang, J., Shah, D., Chen, X., Anderson, R.R. & Wu, M.X. A micro-sterile inflammation array as an adjuvant for influenza vaccines. Nat Commun 5, 4447 (2014).

54. Zhang, L. et al. A proof of concept for neutralizing antibody-guided vaccine design against SARS-CoV-2. National Science Review 8 (2021).

55. Sui, Y. et al. Protection against SARS-CoV-2 infection by a mucosal vaccine in rhesus macaques. JCI Insight 6 (2021).

56. Du, Y. et al. Intranasal administration of a recombinant RBD vaccine induced protective immunity against SARS-CoV-2 in mouse. Vaccine 39, 2280–2287 (2021).

57. An, X. et al. Single-dose intranasal vaccination elicits systemic and mucosal immunity against SARS-CoV-2. iScience 24, 103037 (2021).

58. Bolles, M. et al. A Double-Inactivated Severe Acute Respiratory Syndrome Coronavirus Vaccine Provides Incomplete Protection in Mice and Induces Increased Eosinophilic Proinflammatory Pulmonary Response upon Challenge. Journal of Virology 85, 12201–12215 (2011).

59. Rosati, M. et al. Increased immune responses in rhesus macaques by DNA vaccination combined with electroporation. Vaccine 26, 5223–5229 (2008).

60. Zhao, J. et al. Airway Memory CD4+ T Cells Mediate Protective Immunity against Emerging Respiratory Coronaviruses. Immunity 44, 1379–1391 (2016).

61. Mueller, S.N. & Mackay, L.K. Tissue-resident memory T cells: local specialists in immune defence. Nat Rev Immunol 16, 79–89 (2016).

62. Cohn, B.A., Cirillo, P.M., Murphy, C.C., Krigbaum, N.Y. & Wallace, A.W. SARS-CoV-2 vaccine protection and deaths among US veterans during 2021. Science 0, eabm0620 (2021).

63. Mutsch, M. et al. Use of the inactivated intranasal influenza vaccine and the risk of Bell’s palsy in Switzerland. N Engl J Med 350, 896–903 (2004).

64. Izurieta, H.S. et al. Adverse events reported following live, cold-adapted, intranasal influenza vaccine. JAMA 294, 2720–2725 (2005).

65. Birchall, J.C., Kellaway, I.W. & Gumbleton, M. Physical stability and in-vitro gene expression efficiency of nebulised lipid–peptide–DNA complexes. International Journal of Pharmaceutics 197, 221–231 (2000).

66. Guan, S., Darmstadter, M., Xu, C. & Rosenecker, J. In Vitro Investigations on Optimizing and Nebulization of IVT-mRNA Formulations for Potential Pulmonary-Based Alpha-1-Antitrypsin Deficiency Treatment. Pharmaceutics 13 (2021).

