## Supplementary Information for "Respiratory mucosal vaccination of peptide-poloxamine-DNA nanoparticles provides complete protection against lethal SARS-CoV-2 challenge"

**This file includes:**

Supplementary Table 1 and 2

Supplementary Fig. 1 to 22

Supplementary Results

Supplementary References

Supplementary Methods

Supplementary Notes

### SUPPLEMENTARY TABLES

**Supplementary Table 1. The structure of the multi-modular peptide used in current study.**

| Sequence |
|---|
| KETWWETWWTEWWTEW**KKKKRRRRRKKKK**GAC*SERSMNF*CG |

The Sequence given in italics represents targeting moiety (ICAM-1/CD54 ligand). Underlined sequence represents anchor moiety used to interact with the hydrophobic blocks poloxamine 704. The sequence given in bold represents cationic moiety that is able to condense nucleic acids and facilitate endosomal escape.

**Supplementary Table 2. Physical characterization (size and zeta potential) of pDNA/PP-sNp prepared by different methods.**

| | Size (nm) | PDI | Zeta (mV) |
|---|---|---|---|
| **Optimized** | 116.6±1.7 | 0.183 | -34.7±0.5 |
| **Non-optimized** | 160.3±1.6 | 0.387 | -33.8±0.3 |

The amount of all components used to prepare the nanoparticles were applied according to the “in vivo applications” setting described in “METHODS”. The results are expressed as mean ± SEM of three independent experiments.

### SUPPLEMENTARY FIGURES

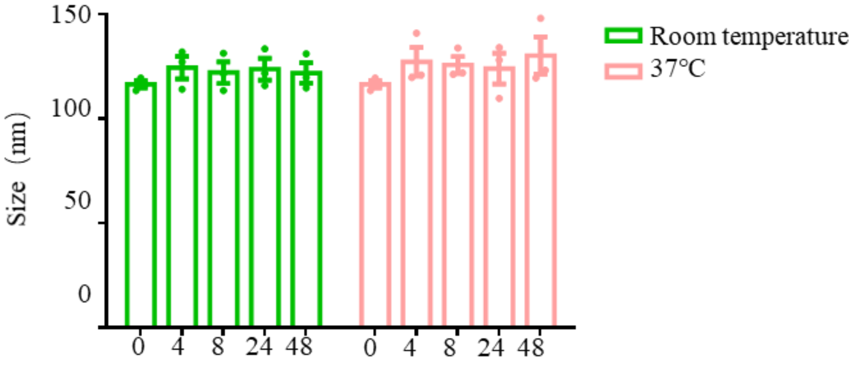


**Supplementary Fig. 1 | Supplementary investigations on the stability of pDNA/PP-sNp.** The hydrodynamic diameters of pDNA/PP-sNp at the indicated time points and temperatures. Data represent the mean ± SEM (n = 3 independent tests)

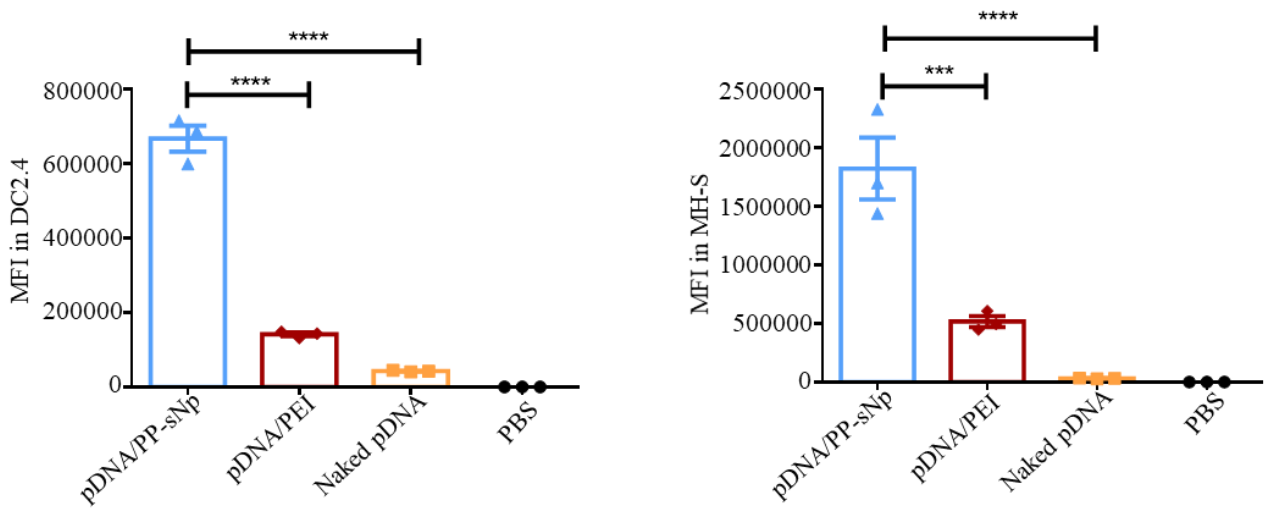


**Supplementary Fig. 2 | Quantitative investigations on the cellular uptake of Cy5-pDNA based formulations via flow cytometry analysis.** Cellular uptake of Cy5-pDNA/PP-sNp, Cy5-pDNA/PEI and naked Cy5-pDNA in DC2.4 (left) and MH-S cells (right). Cy5-fluorescence intensity within the cells was measured by flow cytometry. Data represent the mean ± SEM of three independent experiments. One-way ANOVA with Dunnett's multiple comparisons tests was used to determine significance (****P*<0.001, *****P*<0.0001).

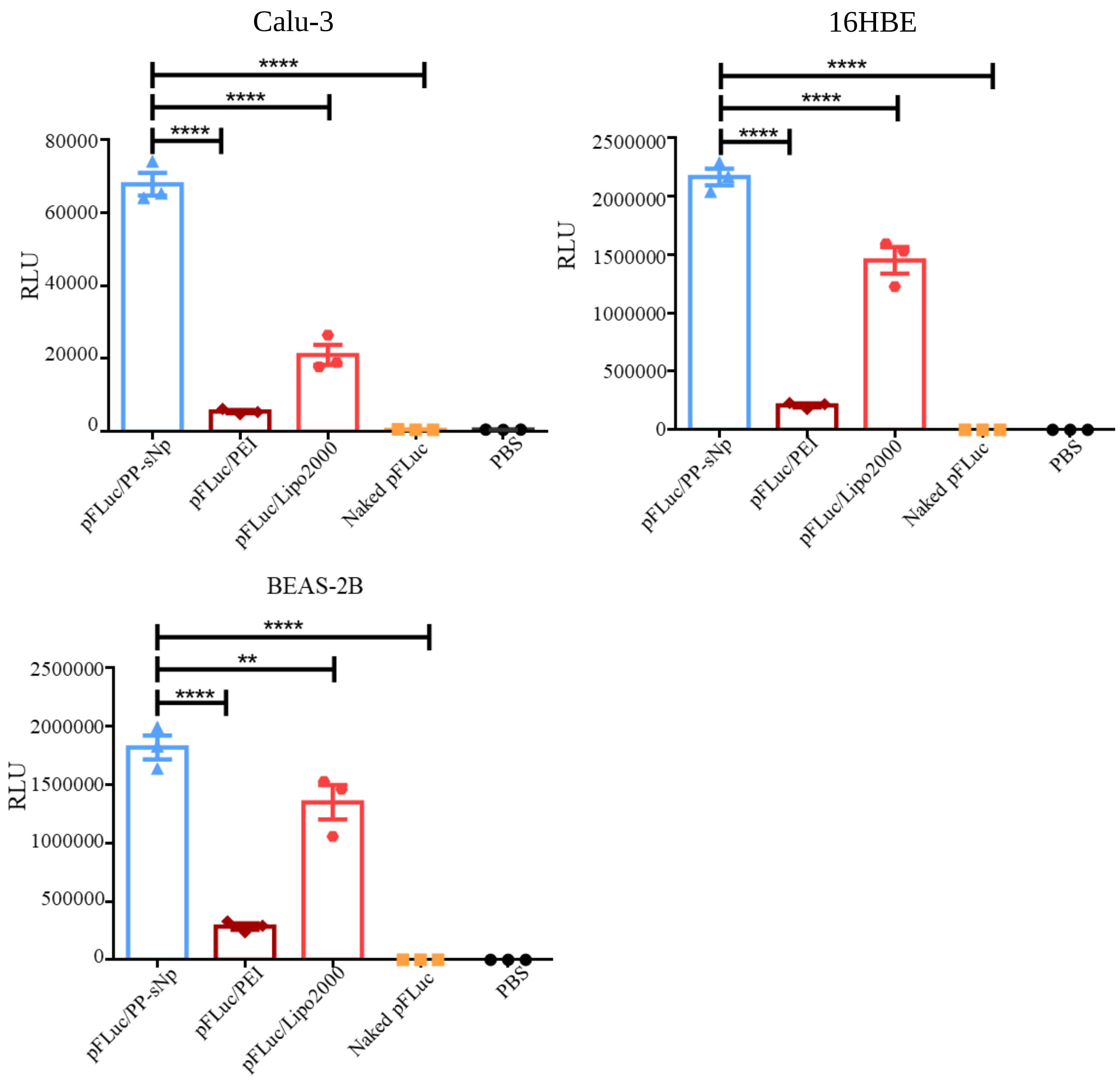


**Supplementary Fig. 3 | Supplementary in vitro transfection efficiency of pFLuc/PP-sNp and control formulations in Calu-3, 16HBE and BEAS-2B cells.** pFLuc/PP-sNp, pFLuc/PEI, pFLuc/Lipo2000 (Lipo2000 represents Lipofectamine 2000) or naked pFLuc were incubated with Calu-3 (upper left), 16HBE (upper right) and BEAS-2B (lower left) cells for 4 h, respectively. Then subjected to the detection of bioluminescence in cell lysate 48 h post-transfection. Data are shown as the mean ± SEM of three independent experiments. Statistical significance was calculated by one-way ANOVA with Dunnett's multiple comparisons tests (***P*<0.01, *****P*<0.0001).

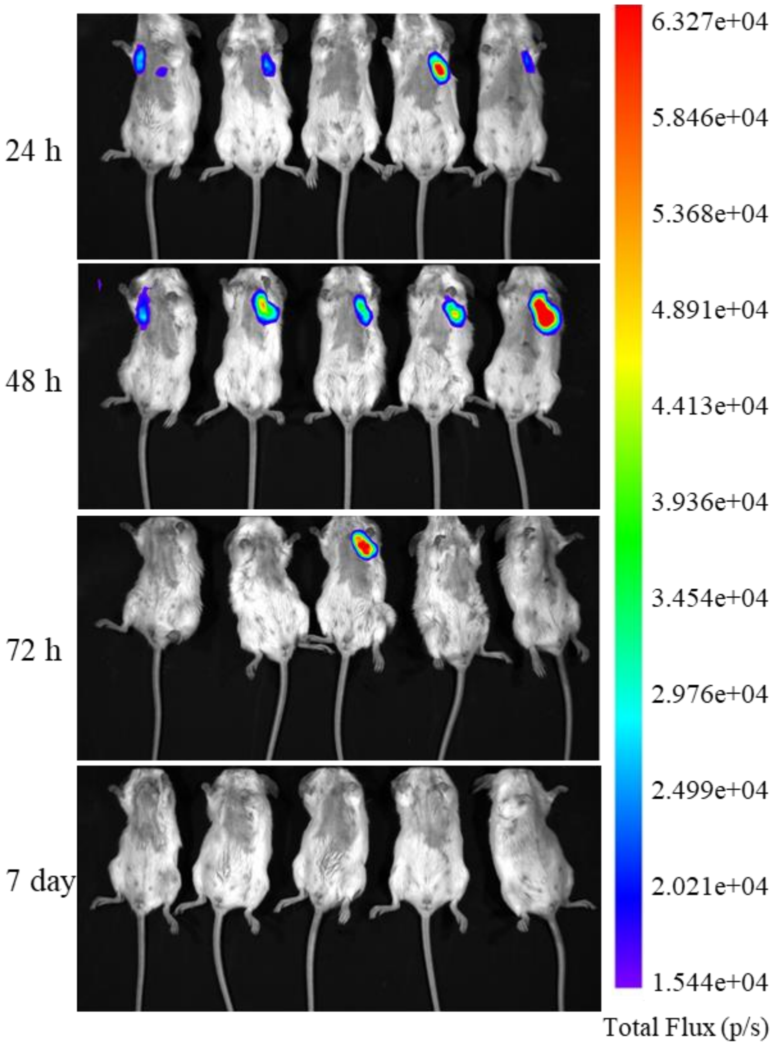


**Supplementary Fig. 4 | Supplementary images of bioluminescence mediated by intratracheally administered pFLuc/PP-sNp at indicated time points in live mice**.

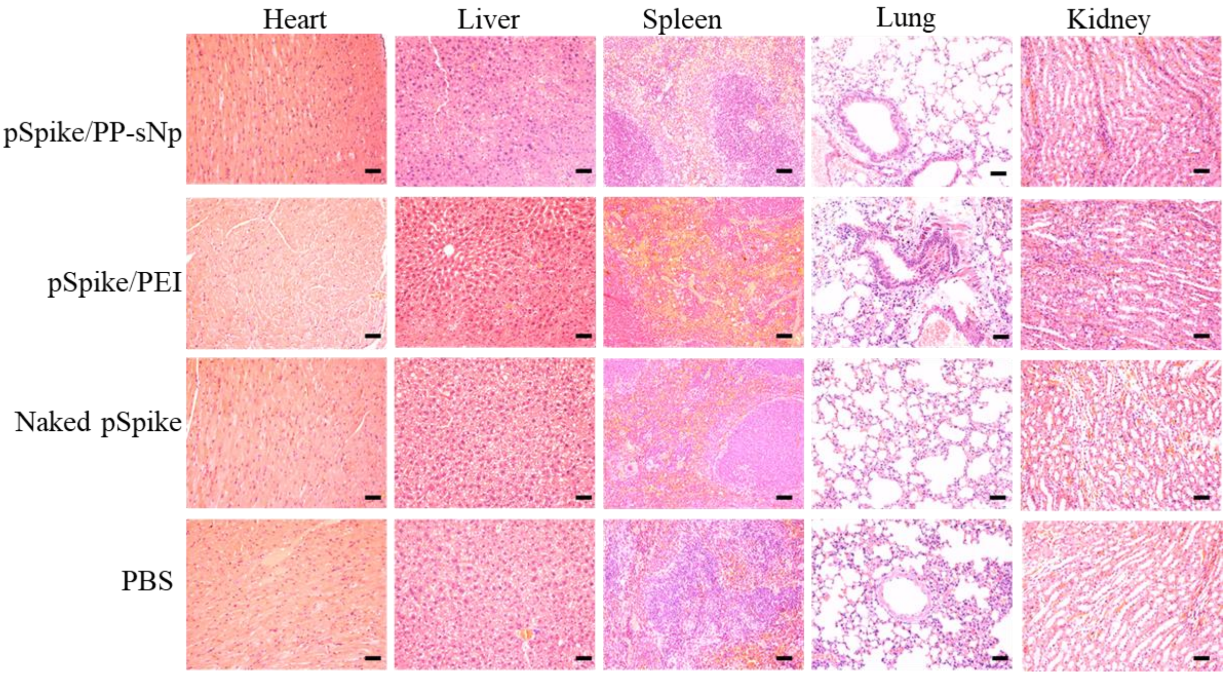


**Supplementary Fig. 5 | Pathohistological analysis of tissue slides from mice in order to evaluate the biocompatibility of pSpike/PP-sNp and control formulations.** Representative hematoxylin and eosin (H&E) staining of organs sections (including the lung, the heart, the liver, the spleen and the kidney) of mice collected 2 days after intratracheal administration of pSpike/PP-sNp, pSpike/PEI or naked pSpike. Control mice were treated with phosphate buffered saline (PBS). Scale bars: 100 μm.

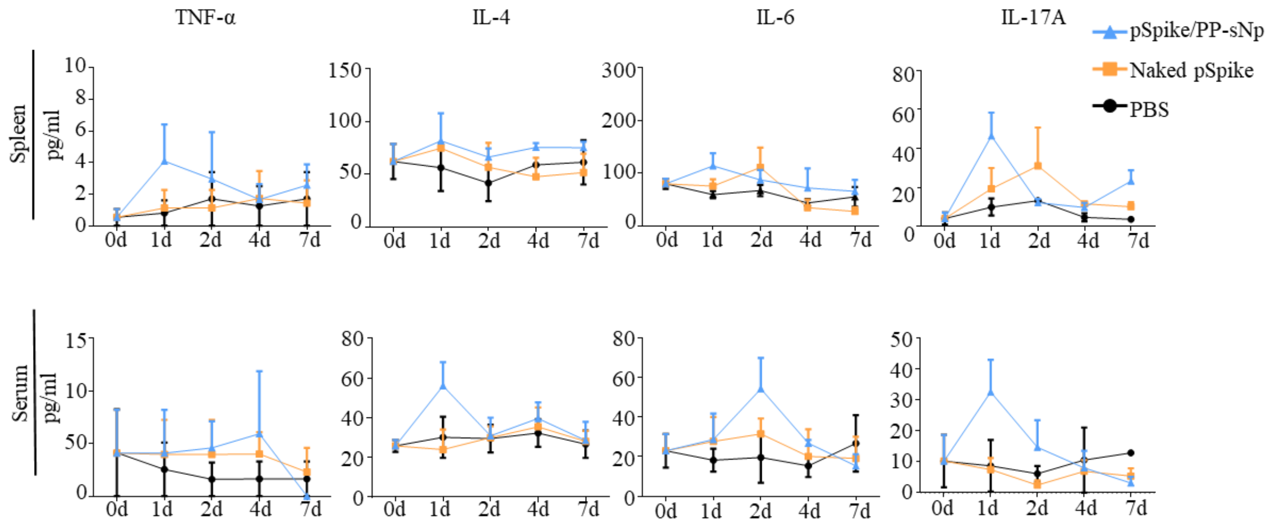


**Supplementary Fig. 6 | Cytokine levels detected in the supernatants of spleen homogenates and serum samples.** The samples were collected from mice intratracheally treated by pSpike/PP-sNp, naked-pSpike or PBS at indicated time points and were analyzed via enzyme-linked immunosorbent assay (ELISA). Data are shown as mean ± SEM. One-way ANOVA with Dunnett's multiple comparisons tests was used to determine significance. No significant difference was detected between groups.

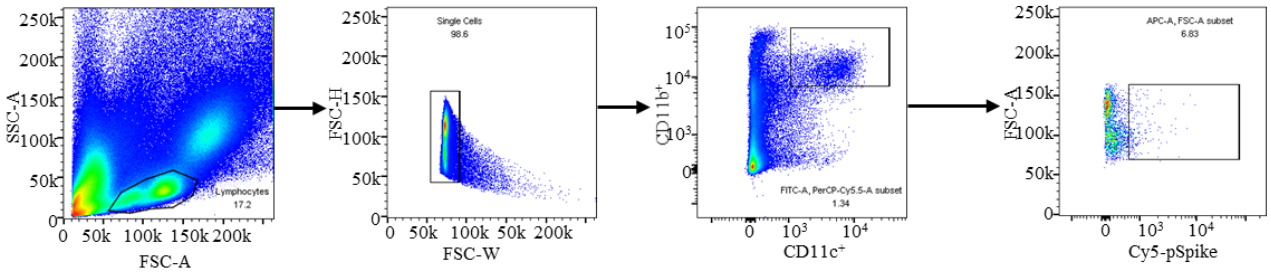


**Supplementary Fig. 7 | Gating strategy used for the flow cytometry analyses as shown in Figure 2e.**

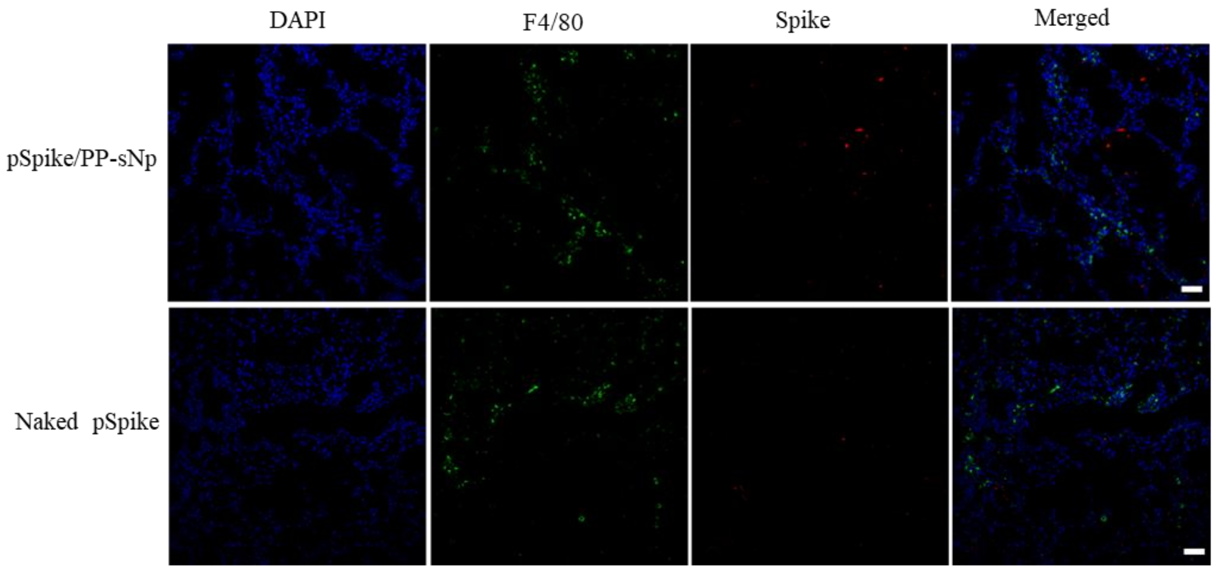


**Supplementary Fig. 8 | Immunofluorescence of lung sections from mice treated by pSpike/PP-sNp and naked pSpike.** The expression of spike protein (Red) and F4/80 (Green). Cell nuclei are stained with DAPI (blue), scale bars: 25 μm.

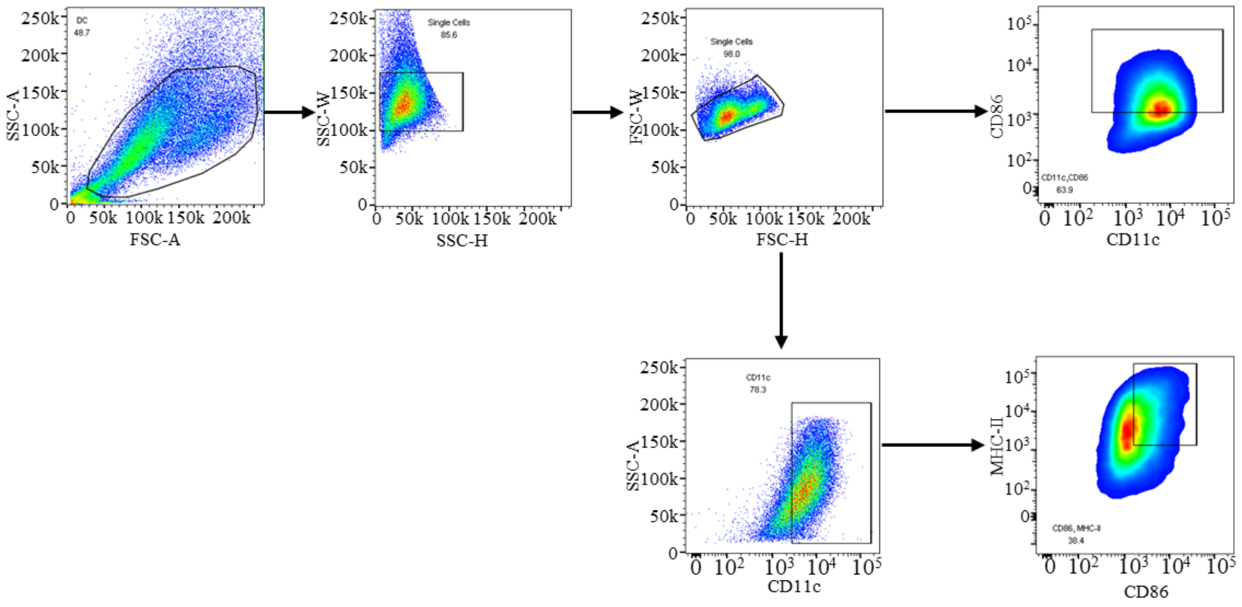


**Supplementary Fig. 9 | Gating strategy used for the flow cytometry analyses as shown in Figure 2h and 2i.**

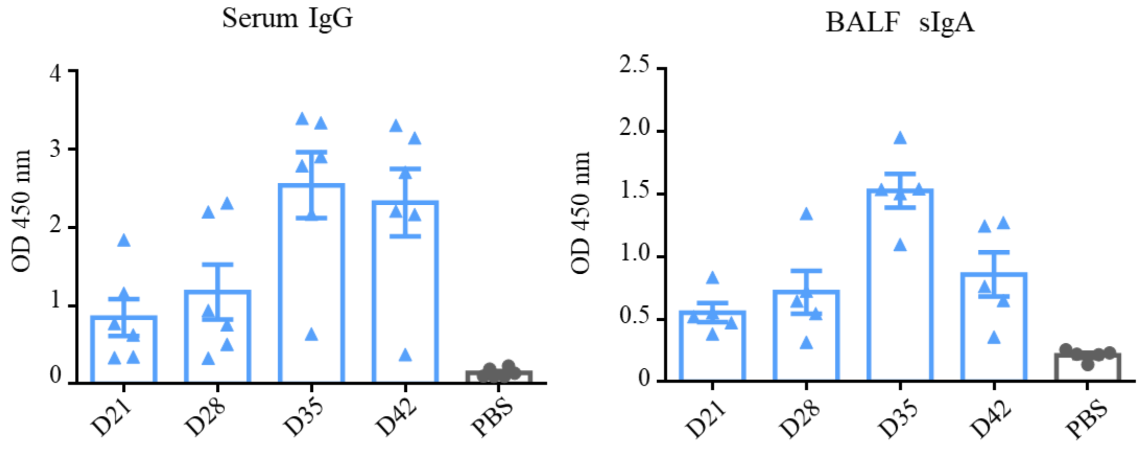


**Supplementary Fig. 10 | The detection of IgG in serum and sIgA in broncho alveolar lavage fluid (BALF) samples of mice vaccinated by pSpike/PP-sNp at different time points of the immunization scheme.** The serum samples were measured with a dilution of 1:50. The BALF samples were measured with no dilution. PBS treated group was used as a control. Data represent the mean ± SEM (n = 5 biologically independent samples).

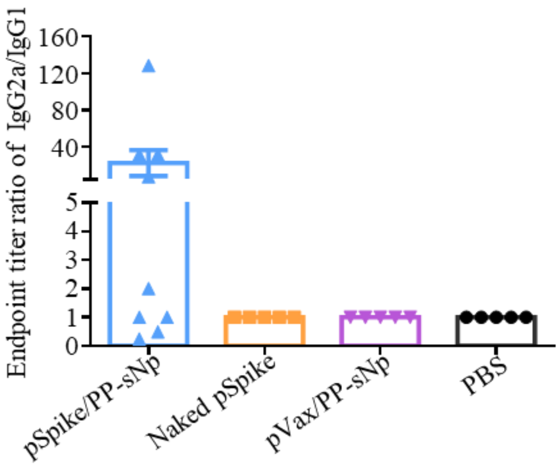


**Supplementary Fig. 11 | The ratio of serum IgG2a to IgG1 (IgG2a/IgG1) binding endpoint titers against SARS-CoV-2 spike (S1 + S2) protein on day 35 of the immunization scheme.** Serum samples were collected from mice intratracheally treated by pSpike/PP-sNp, naked pSpike, pVax/PP-sNp and PBS. Each symbol represents one sample from a biologically independent mouse.

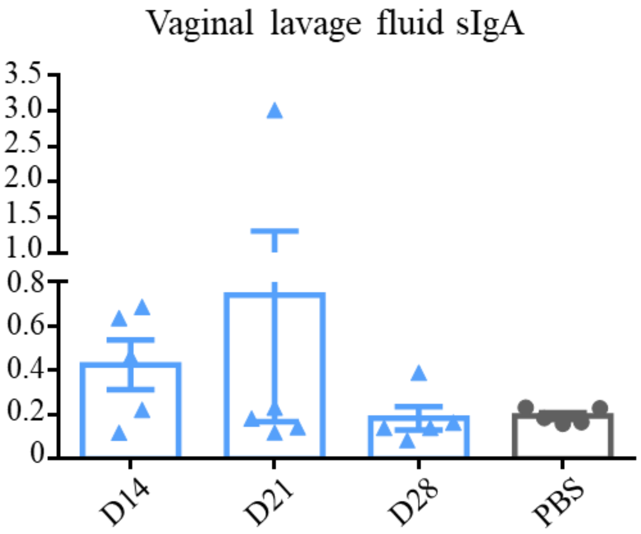


**Supplementary Fig. 12 | The detection of sIgA in vaginal lavage fluid samples collected from pSpike/PP-sNp vaccinated mice at different time points of the immunization scheme**. The samples were measured without further dilution. Samples from PBS treated mice was used as controls. Data represent the mean ± SEM (n = 5 biologically independent samples).

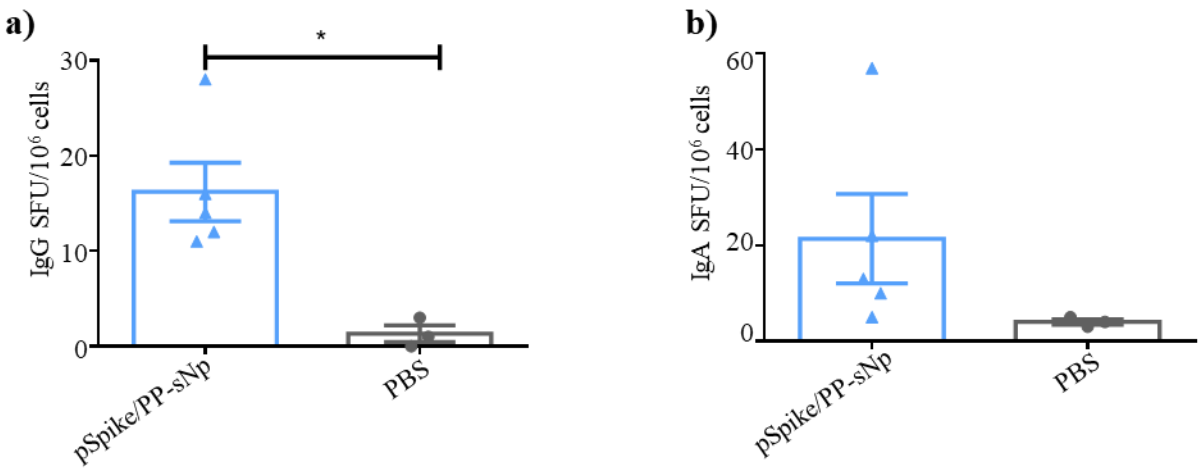


**Supplementary Fig. 13 | The frequency of SARS-CoV-2 spike (S1 + S2) specific a) IgG$^+$ and b) IgA$^+$ antibody secreting cells in the spleen samples collected from pSpike/PP-sNp vaccinated mice via pulmonary route.** The samples were collected 1 week post the 2$^{nd}$ boost, and were measured by an enzyme-linked immunospot (ELISpot) assay. PBS treated group was used as a control. Data represent the mean ± SEM, each symbol represents one sample from a biologically independent mouse. Statistical significance was calculated by a two-tailed unpaired *t*-test (*$P$<0.05).

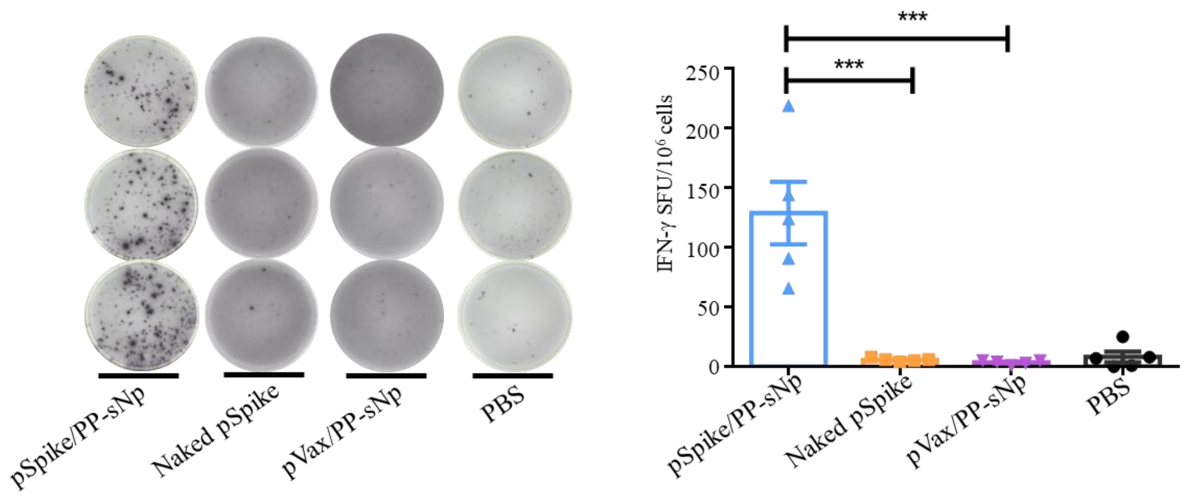


**Supplementary Fig. 14 | ELISpot analysis of IFN-γ spot-forming cells in splenocyte.** The splenocyte was collected from pSpike/PP-sNp, naked pSpike, pVax/PP-sNp or PBS treated mice via pulmonary route. The samples were subjected to ex-vivo stimulation with pools of 14-mer overlapping peptides spanning the SARS-CoV-2 receptor binding domain (RBD) protein. Data represent the mean ± SEM (n = 5 biologically independent samples). One-way ANOVA with Dunnett's post-hoc test was used to determine significance (***P< 0.001).

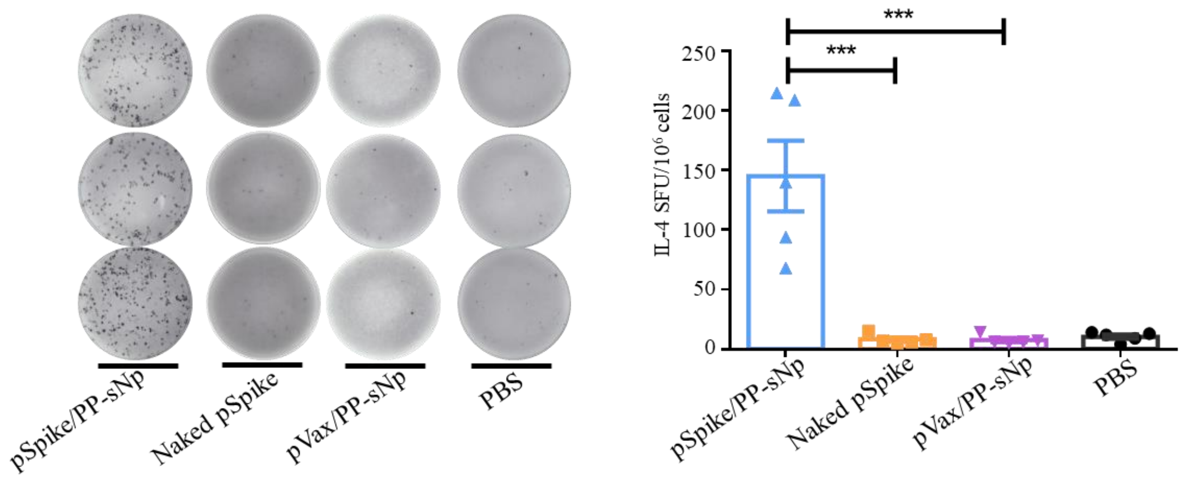


**Supplementary Fig. 15 | ELISpot assay for IL-4 spot-forming cells in splenocyte.** The splenocyte was collected from pSpike/PP-sNp, naked pSpike, pVax/PP-sNp or PBS treated mice via pulmonary route. The samples were subjected to ex-vivo stimulation with pools of 14-mer overlapping peptides spanning the SARS-CoV-2 RBD protein. Data represent the mean ± SEM (n = 5 biologically independent samples). One-way ANOVA with Dunnett's post-hoc test was used to determine significance (****P*< 0.001).

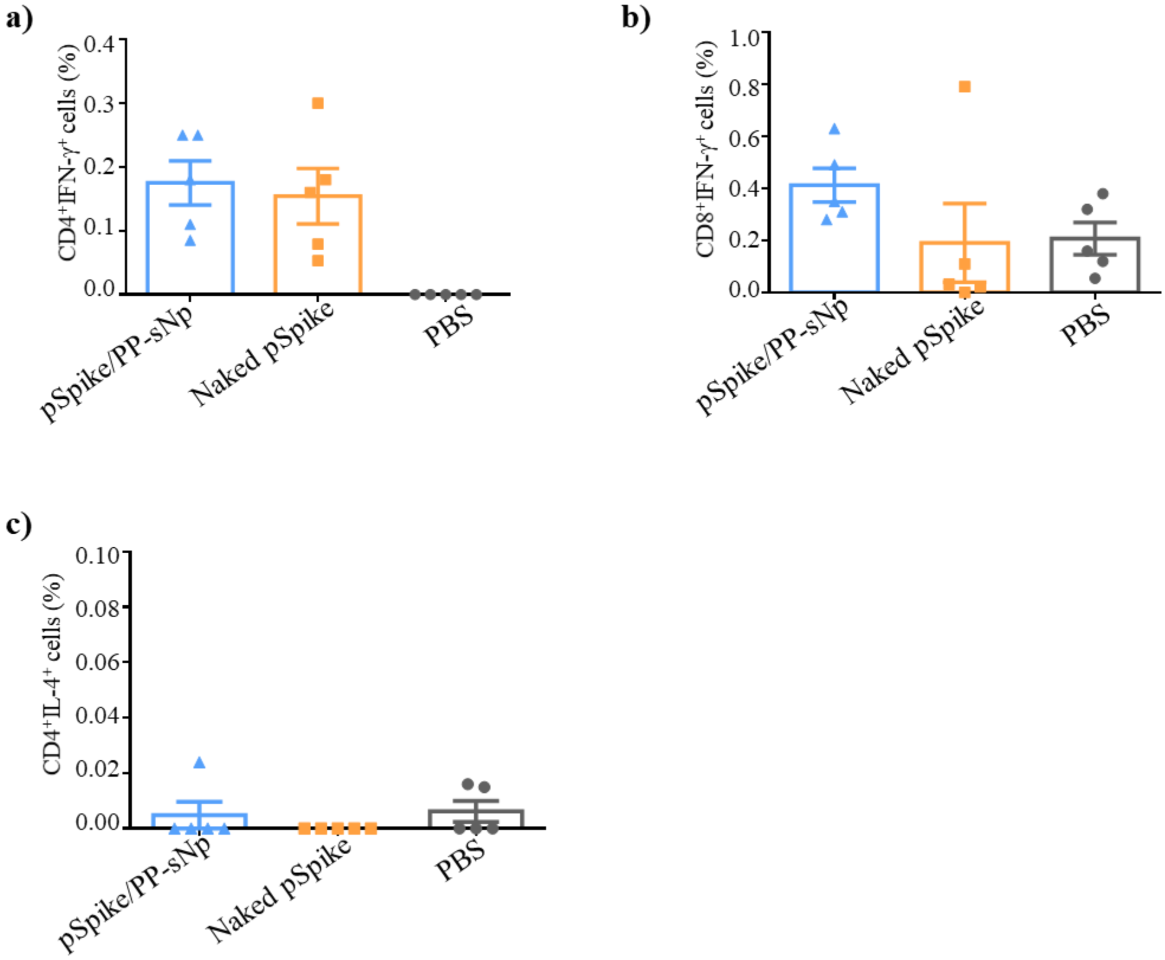


**Supplementary Fig. 16 | Flow cytometry analysis of intracellular cytokines in $CD4^+$ or $CD8^+$ T cells isolated from the spleen of mice.** **a**) $CD4^+$ T cells and **b**) $CD8^+$ T cells in splenocyte of mice intratracheally treated by pSpike/PP-sNp, naked pSpike or PBS were assayed for IFN-$\gamma^+$ expression by flow cytometry after re-stimulation with peptide pools of 14-mer overlapping peptides spanning the SARS-CoV-2 RBD protein. **c**) $CD4^+$ T cells in the spleen of mice were analyzed for IL-$4^+$ expression using the same way as describe above. Data are shown as mean ± SEM of five biologically independent samples and analyzed by one-way ANOVA with Dunnett's post-hoc test to determine the significance. No significant difference was detected between groups.

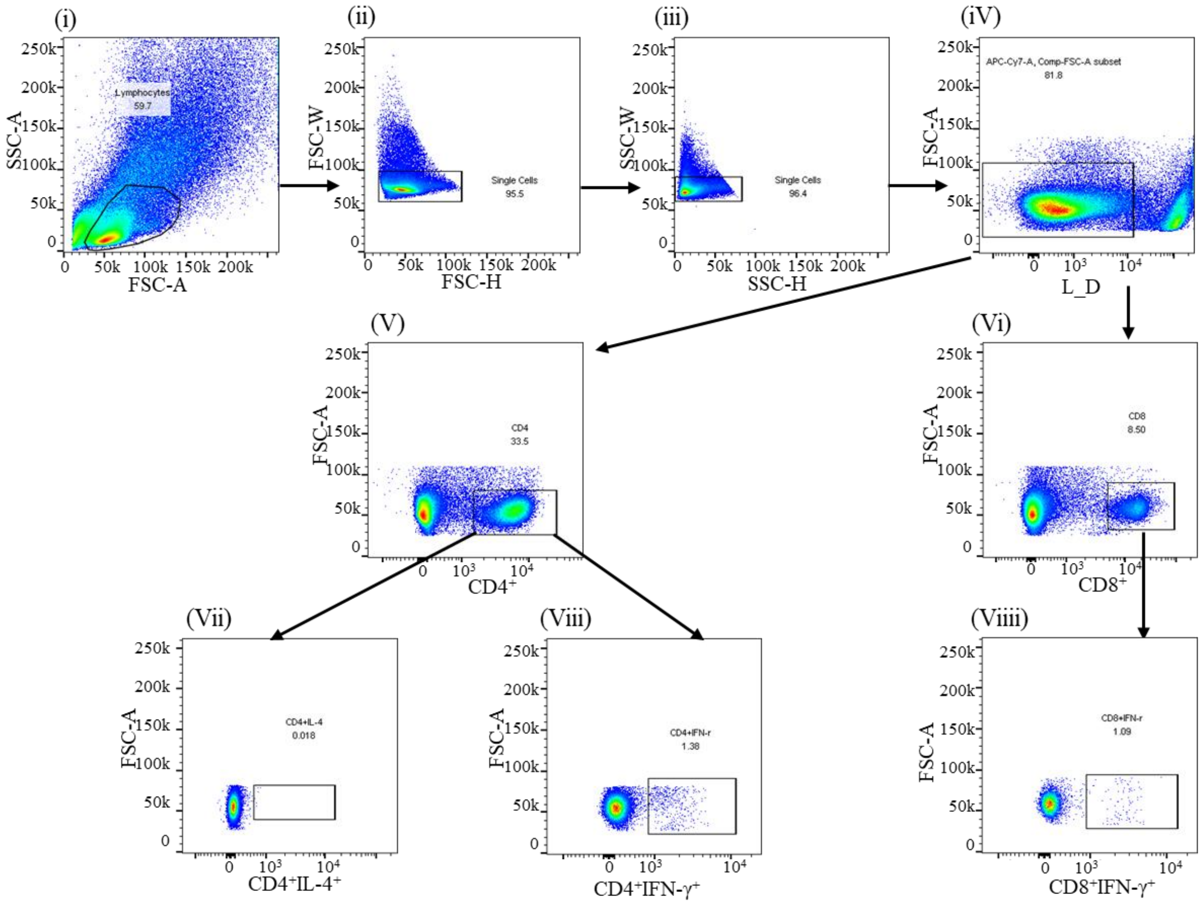


**Supplementary Fig. 17 | Gating strategy used for the flow cytometric analysis of IFN-γ and IL-4 upon SARS-CoV-2 RBD peptide pools stimulation cell populations.** Lymphocytes were gated on (i), then singlets (ii) and (iii) followed by live cells (iV). Next, $CD4^+$ (V) and $CD8^+$ cells were gated (Vi) and from that population $CD4^+IL\text{-}4^+$(Vii), $CD4^+IFN\text{-}\gamma^+$ (Viii) and $CD8^+IFN\text{-}\gamma^+$ (Viiii) T-cells were gated.

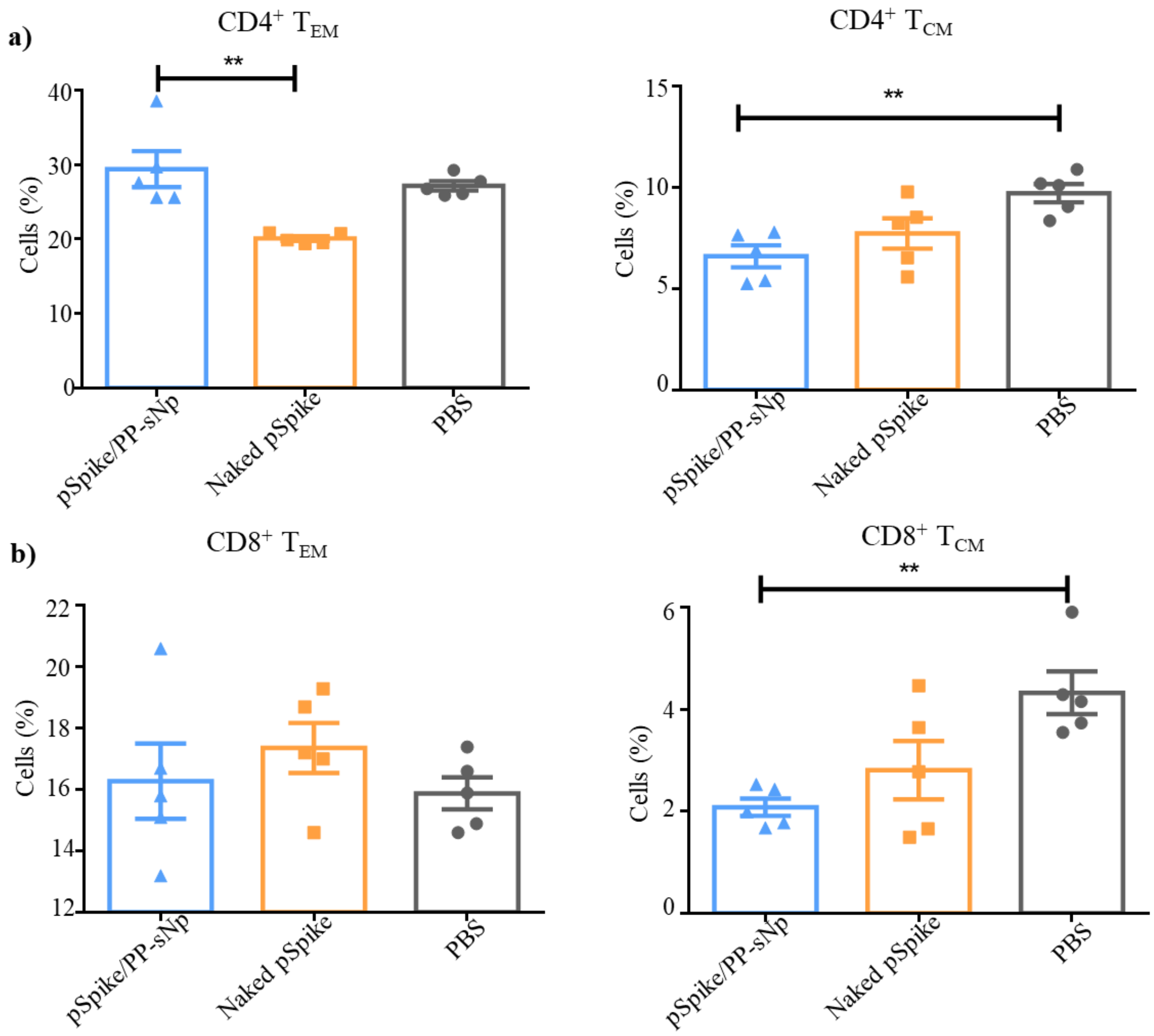


**Supplementary Fig. 18 | Percentage of $T_{EM}$ and $T_{CM}$ in splenocyte of pSpike/PP-sNp, naked pSpike, pVax/PP-sNp or PBS treated mice via pulmonary route. a**) Percentage of $CD4^+$ $T_{EM}$ and $T_{CM}$ in spleen of mice and **b**) $CD8^+$ $T_{EM}$ and $T_{CM}$ in spleen of mice. Data represent the mean ± SEM (n = 5 biologically independent samples). All samples were collected on day 35 of the immunization scheme. One-way ANOVA with Dunnett's post-hoc test was used to determine significance (**$P$< 0.01).

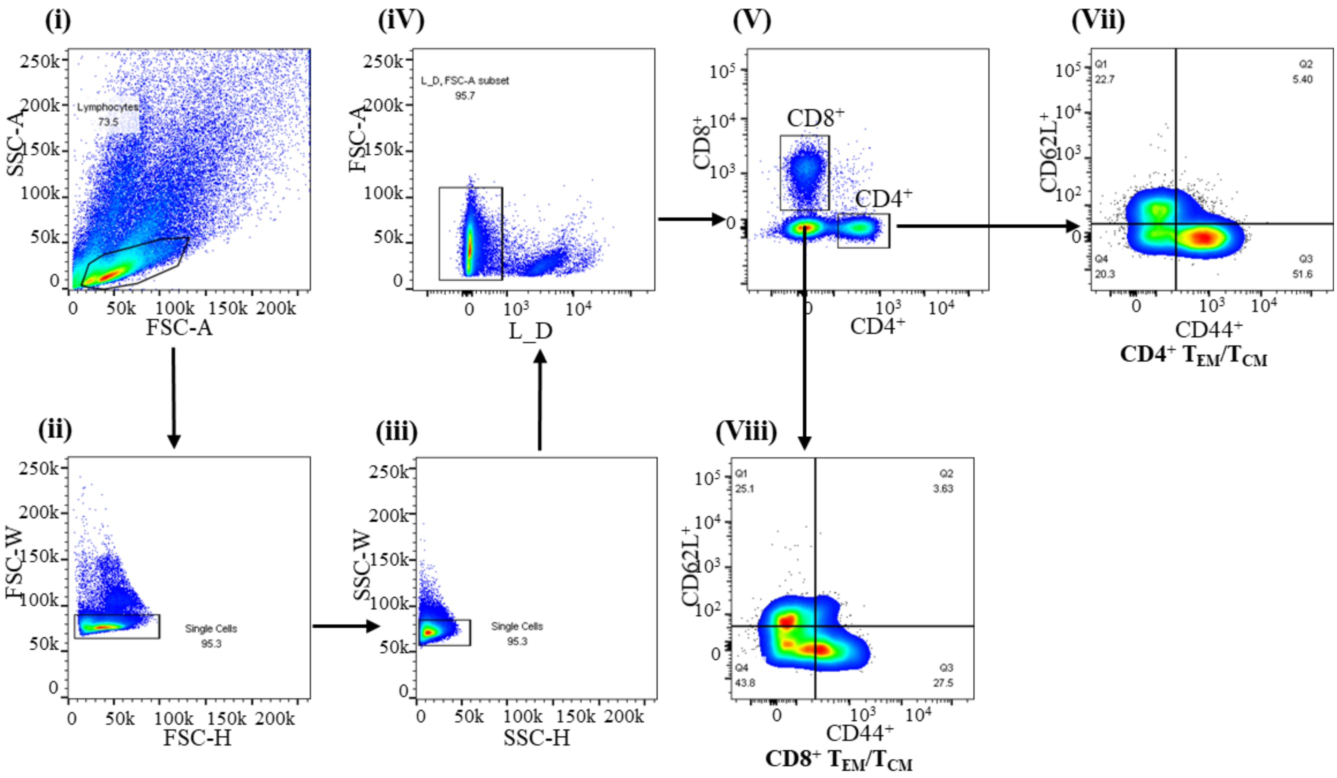


**Supplementary Fig. 19 | Gating strategy used for the flow cytometric analysis of $T_{EM}/T_{CM}$ cell populations.** Lymphocytes were gated on (i), then singlets (ii) and (iii) followed by live cells (iV). Next, $CD4^+$ and $CD8^+$ cells were gated (V) and from that population $CD4^+$ $T_{EM}$ (Vii) and $CD8^+$ $T_{EM}$ (Viii) T-cells were gated.

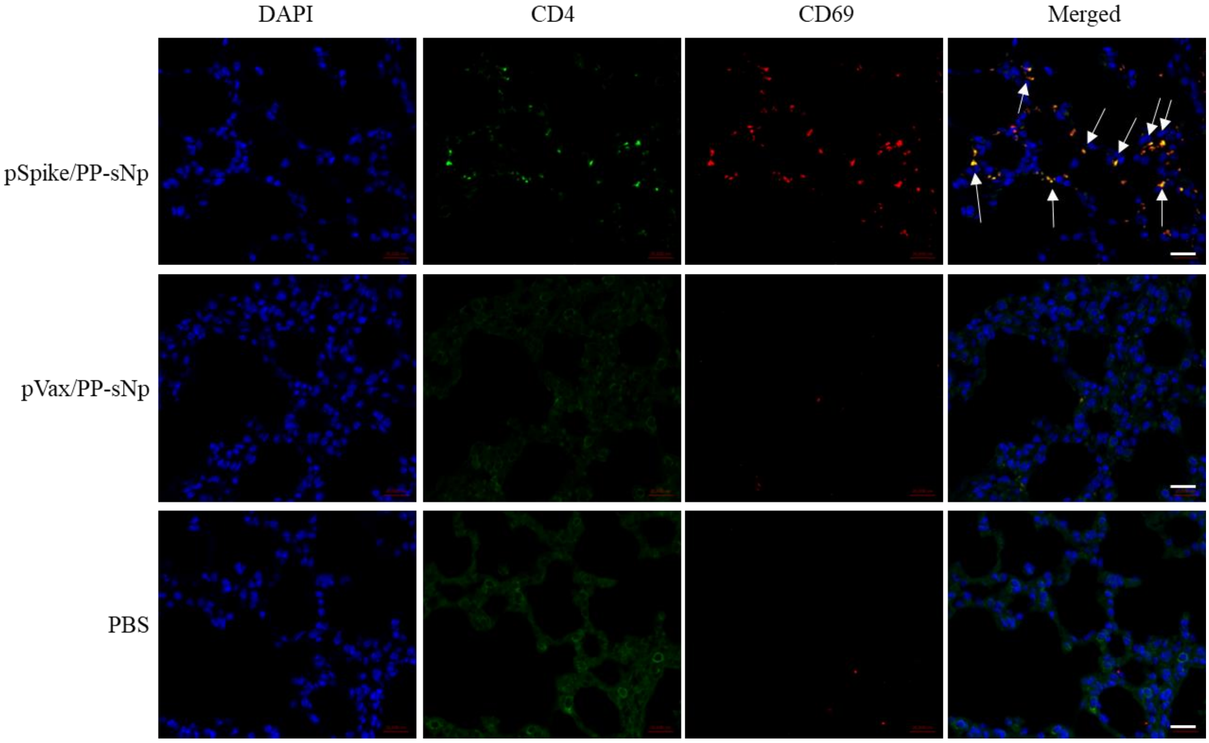


**Supplementary Fig. 20 | Confocal images of $CD4^+$ $T_{RM}$ in lung sections of mice after the SARS-CoV-2 C57MA14 strain challenge.** The samples were collected from pSpike/PP-sNp, pVax/PP-sNp or PBS treated mice 3 days post-challenge. Blue: nucleus, green: CD4, red: CD69, scale bar: 20 μm.

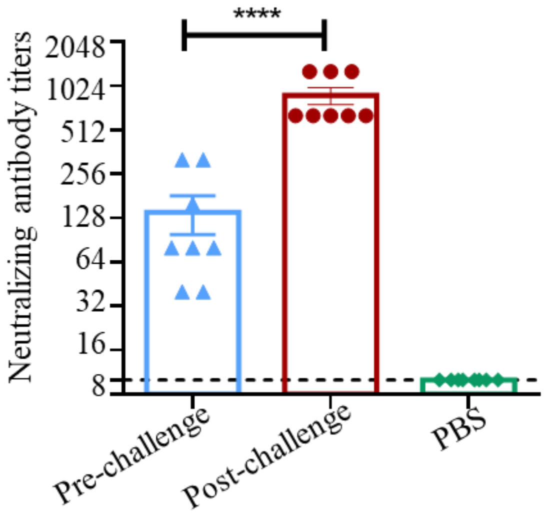


**Supplementary Fig. 21 | The neutralizing antibody titers against SARS-CoV-2 C57MA14 strain in serum of pSpike/PP-sNp immunized mice before and after challenge.** Samples from PBS treated mice was used as controls. Data represent the mean ± SEM (n = 8 biologically independent samples). One-way ANOVA with Dunnett's post-hoc test was used to determine significance (*****P*<0.0001).

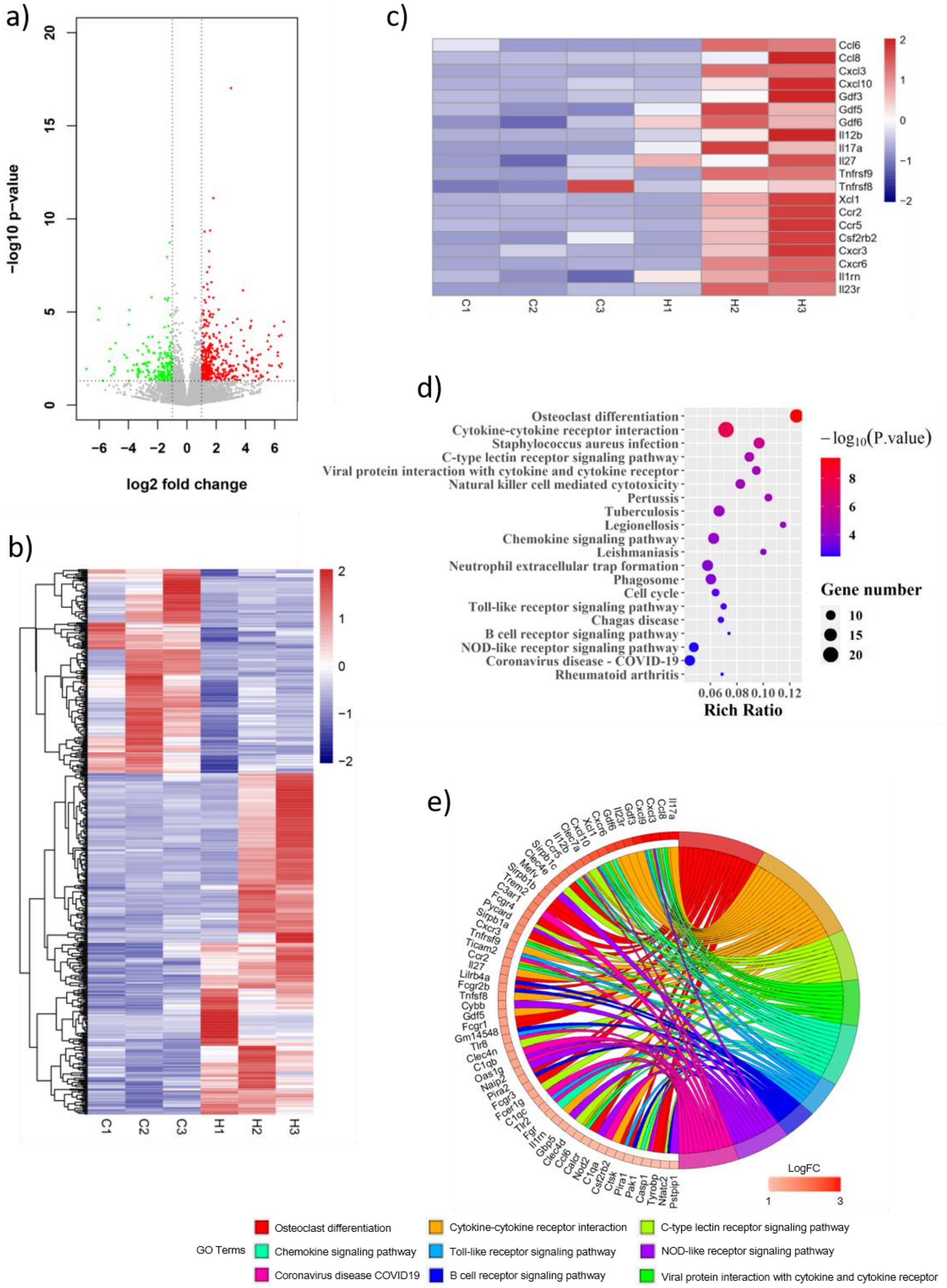


**Supplementary Fig. 22 | RNAseq analysis of mouse lung tissues immunized with pSpike/PP-sNp. a**) Volcano plot and **b**) hierarchical clustering of differentially expressed genes ($P < 0.05$) in lung tissues treated with pSpike/PP-sNp, in comparison to genes expressed in lung tissues of mice treated by PBS. Downregulated genes (green or blue); upregulated genes (red)

**c**) KEGG enrichment analysis of upregulated genes. Shown are the top 20 statistically significant pathways, calculated using an adjusted $P$ value < 0.05. **d**) Heat map representing the upregulated cytokines, chemokines and receptors. **e**) Chord plot depicting the relationship between PP-sNp-mediated mouse pulmonary immunization-related genes and KEGG pathways.

### SUPPLEMENTARY RESULTS

**PP-sNp induces innate and adaptive antiviral pathways in the lungs of mice**

To further understand the function of pSpike/PP-sNp in pulmonary immunization, we performed RNA sequencing of lung tissues of pSpike/PP-sNp immunized mouse, with PBS-treated mice were used as control. A total of 623 differentially expressed genes (DEGs) were identified, including 427 up-regulated genes and 196 down-regulated genes (Supplementary Fig. 22a and 22b). A variety of cytokines (IL-12b, IL-17a, IL-27) and chemokines (CCL6, CCL8, CXCL3, CXCL9, CXCL10), which involved in immunoregulatory and inflammatory processes, were detected among the up-regulated genes (Supplementary Fig. 22c). KEGG enrichment analysis showed that the up-regulated genes were significantly enriched in pro-inflammatory signalling (e.g. Cytokine-cytokine receptor interaction, Chemokine signalling pathway, Toll-like receptor signalling pathway and NOD-like receptor signalling pathway). Meanwhile, C-type lectin receptor (CLR) signalling pathway, which induce the expression of specific cytokines to determine T cell polarization fates[1], was significantly enriched as well (Supplementary Fig. 22d). Notably, antiviral defence pathways, including viral protein interaction with cytokine and cytokine receptor and coronavirus disease-COVID-19, also participated in the PP-sNp induced pulmonary immunization (Supplementary Fig. 22d). Multiple genes, such as *NOD2*, *CLEC7A* (also known as *Dectin1*), *IL-17a*, have been reported to participate in innate and adaptive immune response[2–5]. We also found these genes are actively involved in the PP-sNp induced pulmonary immunization through a variety of pathways (Supplementary Fig. 22e).

The innate immune system play a key role in the recognition and early response to infectious agents. Innate immune cells, such as macrophages and dendritic cells (DCs), recognize viruses through extra- and intracellular receptors (including Toll-like and NOD-like receptors) and secrete inflammatory mediators (including chemokines and cytokines)[6], which enhance antigen processing and in turn active T and B cells[7]. We found that Toll-like receptor and NOD-like receptor signalling pathways were significantly enriched in pSpike/PP-sNp induced pulmonary immunization. Three key mediator genes, *TLR2*, *TLR8* and *NOD2*, which were associated with pathogen-mediated inflammatory response to innate immunity[2,8], were significantly upregulated in lung tissues of pSpike/PP-sNp immunized mouse, indicating that PP-sNp can induce innate immune response. C-type lectin receptors (CLRs) expressed by dendritic cells are crucial for tailoring immune responses to pathogens and CLR signalling pathway play a key role in adaptive immune responses. Most CLRs induce adaptive immune responses via activate canonical NF-κB pathway, however, *CLEC7A* induce an unique signalling pathway that leads to the activation of the non-canonical NF-κB pathway[1,3]. Collectively, we inferred that PP-sNp mediated innate and adaptive immune responses through the recognition by pattern recognition receptors (PRRs) of pulmonary immune cells, but the molecular mechanism needs to be further verified in future studies.

### SUPPLEMENTARY REFERENCES


1. Geijtenbeek, T. B. H. & Gringhuis, S. I. Signalling through C-type lectin receptors: shaping immune responses. *Nat. Rev. Immunol.* **9**, 465–479 (2009).
2. Kim, D. *et al.* Nod2-mediated recognition of the microbiota is critical for mucosal adjuvant activity of cholera toxin. *Nat. Med.* **22**, 524–530 (2016).
3. Saijo, S. *et al.* Dectin-1 is required for host defense against Pneumocystis carinii but not against Candida albicans. *Nat. Immunol.* **8**, 39–46 (2007).
4. Christensen, D., Mortensen, R., Rosenkrands, I., Dietrich, J. & Andersen, P. Vaccine-induced Th17 cells are established as resident memory cells in the lung and promote local IgA responses. *Mucosal Immunol.* **10**, 260–270 (2017).
5. Shibui, A. *et al.* Th17 cell-derived IL-17 is dispensable for B cell antibody production. *Cytokine* **59**, 108–114 (2012).
6. Nachbur, U. *et al.* A RIPK2 inhibitor delays NOD signalling events yet prevents inflammatory cytokine production. *Nat. Commun.* **6**, 6442 (2015).
7. Hoebe, K., Janssen, E. & Beutler, B. The interface between innate and adaptive immunity. *Nat. Immunol.* **5**, 971–974 (2004).
8. Kumar, H., Kawai, T. & Akira, S. Toll-like receptors and innate immunity. *Biochem. Biophys. Res. Commun.* **388**, 621–625 (2009).

### SUPPLEMENTARY METHODS

**RNA sequencing.** Lung tissues were collected 1 week after the last immunization. Total RNAs isolation, cDNA synthesis and the RNA-seq library was prepared by the Beijing Genomics Institute. Sequence reads were obtained using BGISEQ-500 (Illumina) and successfully mapped to mouse genome. Reads counts were normalized on the basis of fragments per kilobase per million (FPKM), fold changes were calculated for all possible comparisons, and a twofold cutoff with an adjusted $P$ value < 0.05 was used to select genes with expression changes. The set of expressed genes were used as for functional enrichment analyses.

**SUPPLEMENTARY NOTES**

**Primer Sequence of SARS-CoV-2 S protein**

**Forward** 5'-ATCGGCATCGTGAACAACAC-3'

**Reverse** 5'-AGCCAGATGTACCAAGGCCA-3'

**Peptide Pools of 14-mer Overlapping Peptides Spanning the SARS-CoV-2 RBD Protein**

| NO | Sequence | NO | Sequence |
|---|---|---|---|
| 1 | RVQPTESIVRFPNI | 23 | FTGCVIAWNSNNLD |
| 2 | ESIVRFPNITNLCP | 24 | IAWNSNNLDSKVGG |
| 3 | FPNITNLCPFGEVF | 25 | NNLDSKVGGNYNYL |
| 4 | NLCPFGEVFNATRF | 26 | KVGGNYNYLYRLFR |
| 5 | GEVFNATRFASVYA | 27 | YNYLYRLFRKSNLK |
| 6 | ATRFASVYAWNRKR | 28 | RLFRKSNLKPFERD |
| 7 | SVYAWNRKRISNCV | 29 | SNLKPFERDISTEI |
| 8 | NRKRISNCVADYSV | 30 | FERDISTEIYQAGS |
| 9 | SNCVADYSVLYNSA | 31 | STEIYQAGSTPCNG |
| 10 | DYSVLYNSASFSTF | 32 | QAGSTPCNGVEGFN |
| 11 | YNSASFSTFKCYGV | 33 | PCNGVEGFNCYFPL |
| 12 | FSTFKCYGVSPTKL | 34 | EGFNCYFPLQSYGF |
| 13 | CYGVSPTKLNDLCF | 35 | YFPLQSYGFQPTNG |
| 14 | PTKLNDLCFTNVYA | 36 | SYGFQPTNGVGYQP |
| 15 | DLCFTNVYADSFVI | 37 | PTNGVGYQPYRVVV |
| 16 | NVYADSFVIRGDEV | 38 | GYQPYRVVVLSFEL |
| 17 | SFVIRGDEVRQIAP | 39 | RVVVLSFELLHAPA |
| 18 | GDEVRQIAPGQTGK | 40 | SFELLHAPATVCGP |
| 19 | QIAPGQTGKIADYN | 41 | HAPATVCGPKKSTN |
| 20 | QTGKIADYNYKLPD | 42 | VCGPKKSTNLVKNK |
| 21 | ADYNYKLPDDFTGC | 43 | KSTNLVKNKCVNF |
| 22 | KLPDDFTGCVIAWN | | |
